# Peripheral immune cell response to stimulation stratifies Parkinson’s disease progression from prodromal to clinical stages

**DOI:** 10.1101/2024.12.05.625499

**Authors:** Julian R. Mark, Ann M. Titus, Hannah A. Staley, Stephan Alvarez, Savanna Mahn, Nikolaus R. McFarland, Rebecca L. Wallings, Malú Gámez Tansey

**Author notes:** Correspondence should be directed to Drs. Malú Gámez Tansey & Rebecca L. Wallings. McKnight Brain Institute, 1149 Newell Dr, Gainesville, FL 32610, United States. Email: Malú Tansey, and Rebecca Wallings.

## Abstract

The motor stage of idiopathic Parkinson’s disease (iPD) can be preceded for years by a prodromal stage characterized by non-motor symptoms like REM sleep behavior disorder (RBD). Here, we show that multiple stages of iPD, including the pre-motor prodromal stage, can be stratified according to the inflammatory and immunometabolic responses to stimulation of peripheral blood mononuclear cells *ex vivo*. We identified increased stimulation-dependent secretion of TNF, IL-1β, and IL-8 in monocytes from RBD patients and showed diminished proinflammatory cytokine secretion in monocytes and T cells in early and moderate stages of PD. Mechanistically, immune activation revealed deficits in CD8^+^ T-cell mitochondrial health in moderate PD, and relative mitochondrial health in CD8^+^ T cells was positively correlated with stimulation-dependent T-cell cytokine secretion across the PD spectrum. Dysregulated immunometabolism may drive peripheral inflammation and PD progression, and *ex vivo* stimulation-based assays have potential to reveal novel biomarkers for patient stratification and progression with immune endophenotypes.

## Main

Parkinson’s disease (PD) is a multi-system neurodegenerative disease for which there are no effective disease modifying therapies. Neuroprotective strategies have been largely ineffective because the majority of dopaminergic neurons in the *substantia nigra pars compacta* (SNpc) have already been lost by the time motor symptoms present and clinical diagnosis can be made^1^. This underscores the need for accessible biomarkers that facilitate early diagnosis and identify patient endophenotypes for superior recruitment and assignment into clinical trials. Significant attention has been directed towards patients with isolated REM sleep behavior disorder (RBD), as this disorder is a strong predictive marker of pre-motor prodromal PD^2, 3^. RBD is characterized by loss of muscle atonia during REM sleep and the physical acting out of dreams that are often intense or violent^4^. Approximately 80% of RBD patients will develop a neurodegenerative disease within 10.5 years of RBD diagnosis, and the plurality (43%) of those who convert will develop PD^2^. Thus, studying individuals with RBD could reveal novel biomarkers for earlier diagnosis of PD and grant insight into the mechanisms which drive disease progression prior to the onset of classical PD-associated motor symptoms.

Dysregulation in the immune system has long been implicated in the pathogenesis of PD^5^, and this has led to the emergence of peripheral immune dysfunction as a promising mechanism with potentially disease-relevant biomarkers. Studies have reported increased tumor necrosis factor (TNF) receptor expression as well as enhanced production of interferon-gamma (IFNγ) and TNF from T cells in PD patients^6^. Furthermore, circulating monocytes from PD patients display upregulation of genes involved in immune activation, including HLA-DQB1, MYD88, REL, and TNF^7^. Together, these findings point towards widespread changes across both the innate and adaptive peripheral immune system in PD. Indeed, it has been shown that the neutrophil-to-lymphocyte ratio (NLR), which serves as a biomarker for systemic inflammation, is significantly correlated with lower levels of dopamine transporter in the striatum of PD patients^8^. Thus, peripheral immune dysregulation may be critical to PD progression, and targeting these mechanisms may improve our ability to deliver personalized treatment plans.

Recent meta-analyses have reported increased blood levels of inflammatory cytokines, such as TNF, IL-1β, and IL-6, in idiopathic PD (iPD) patients compared to controls^9, 10^. It has also been found that carriers of PD-associated mutations in *LRRK2* and *GBA1* exhibit increased serum cytokine levels^11, 12^, highlighting peripheral immune dysfunction as a common theme shared by both idiopathic and genetic forms of PD. However, circulating cytokine levels are subject to significant variability influenced by circadian rhythm, diet, and environmental exposures^13, 14, 15, 16^, which has limited their effectiveness as biomarkers and contributed to heterogeneous reports^17, 18^. Moreover, investigations of plasma cytokine levels in prodromal PD have been inconclusive. One study described increased serum TNF levels in RBD patients^19^, while another reported no differences relative to controls^20^. Therefore, the extent of detectable immune dysfunction in prodromal PD remains to be determined. To facilitate the development of accurate and predictive biomarkers, new approaches are required to overcome background noise with sufficient sensitivity to reveal facets of immune dysfunction that may be difficult to parse apart at baseline.

One approach that our group has demonstrated to help overcome these challenges is to examine differences in immune cell “traits” using stimulus-evoked responses of peripheral immune cells *ex vivo*. These traits are defined by stimulus-evoked activation and resolution responses, and they can be reproducibly elicited in a controlled *ex vivo* experiment regardless of exogenous factors. In contrast, the previous literature on baseline cytokine levels describes immune “states”, reflecting only the content of blood cytokines at a single timepoint when the sample is drawn and which can be highly variable^13, 14, 15, 16, 21, 22^. Epidemiologic studies have linked exposure to environmental pathogens with increased long-term risk for developing PD^23^, and this has raised the possibility that an aberrant immune response to stimulation may be more relevant for predicting PD risk than baseline levels of inflammatory factors in the blood. Stimulation-based assays have the potential to provide greater sensitivity, with α-synuclein peptide exposure shown to elicit increased TNF secretion in lymphocytes from PD patients but not from neurologically healthy controls (NHCs)^24^. Immune stimulation also has the advantage of increasing energetic demand^25^, which can highlight immunometabolic deficits that have been strongly implicated in PD pathogenesis^26, 27, 28^. For example, peripheral blood mononuclear cells (PBMCs) from PD patients show significantly altered mitochondrial respiratory capacity and mitochondrial membrane potential relative to controls^29, 30^, as well as downregulation of a number of lysosome/autophagy related genes including *ULK3*, *ATG2A*, and *HDAC6*^31^. Furthermore, PD monocytes have reduced mitochondrial content relative to controls^32^, and monocyte activity of the lysosomal enzyme glucocerebrosidase is inversely correlated with severity of motor symptoms after diagnosis^33^. Mitochondrial and lysosomal deficits have yet to be reported in PBMCs at the prodromal stage of PD, but it remains possible that metabolic organelle dysfunction is present before motor symptoms manifest yet is too subtle to observe with baseline measurements. It is therefore vital to explore how the stage of PD progression may alter immunometabolic response to activation, as this will not only bolster our understanding of PD etiology but potentially provide novel means of identifying at-risk individuals to recruit into suitable trials and for monitoring disease progression.

To close these important gaps in knowledge, we sought to investigate whether the peripheral immune response to stimulation is dysregulated in RBD patients relative to multiple stages of PD progression. To test this hypothesis, we studied the stimulation-dependent responses of isolated T cells and monocytes from RBD patients, iPD patients at early (within 2 years of diagnosis) and moderate (2-10 years after diagnosis) stage disease, and NHCs. The inclusion of early and moderate iPD groups enabled us to capture the dynamic changes in immunometabolic responses across the disease spectrum. In addition, cell-type specific cytokine secretion was evaluated to enhance our ability to detect immune dysfunction traits and determine for the first time if different PBMC subsets display unique patterns of dysregulation in prodromal versus motor PD. Our results show that RBD patients display a distinct signature of immune activation relative to NHCs and clinically diagnosed PD patients, and immunometabolic function of PBMC subsets enables stratification of PD progression across multiple stages of disease.

## Results

### Monocytes from RBD patients display dysregulated stimulation-dependent cytokine secretion

To determine if PBMC subsets from iPD patients exhibit differences in stimulation-evoked inflammatory cytokine secretion based on disease progression, we began by collecting PBMCs from patients across the disease spectrum. Whole blood samples were collected from RBD patients to approximate prodromal PD (*n* = 15), early-stage PD patients within 2 years of diagnosis (*n* = 27), moderate-stage PD patients within 2-10 years of diagnosis (*n* = 30), and age- and sex-matched neurologically healthy controls (NHC, *n* = 21). Patients were enrolled at the Norman Fixel Institute for Neurological Diseases at the University of Florida (demographic information of the cohorts is shown in Table 1). From whole blood, PBMCs were isolated and cryopreserved using previously published methods^34^. After PBMCs were thawed, CD3^+^ T cells and pan-monocytes were magnetically isolated, plated, and treated with an immune stimulus (200U IFNγ for monocytes and 3.125 μL of CD3/CD28 T-Activator Dynabeads for T cells) or vehicle control for 72 hours (workflow is shown in Fig. 1).

**Fig. 1:**
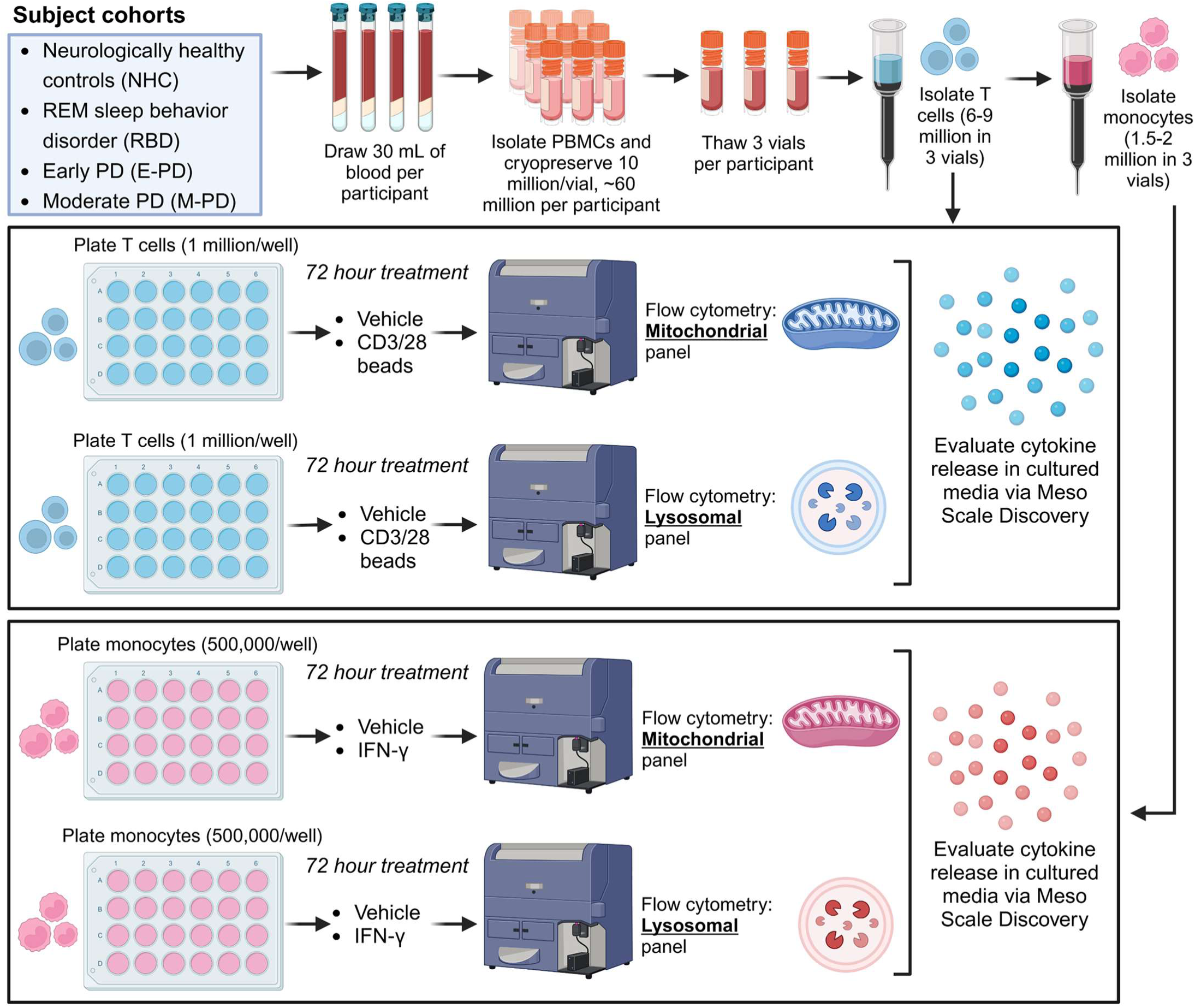
Workflow and experimental design. Whole blood was collected from participants consisting of neurologically healthy controls (NHCs), patients with REM sleep behavior disorder (RBD), early-stage PD patients (diagnosed <2 years prior), and moderate-stage PD patients (diagnosed 2-10 years prior). Peripheral blood mononuclear cells were isolated from whole blood, cryopreserved, and thawed, and subjected to magnetic bead isolations to obtain purified CD3^+^ T cells or purified pan-monocytes. Cells were allowed to rest for 2 hours, followed by 72-hour incubation in presence or absence of a stimulation source (200U IFNγ for monocytes, 3.125 μL of CD3/CD28 Dynabeads for T cells). Then cells were assessed via flow cytometry and media was taken for cytokine quantification. Panel was created with BioRender.com.

**Table 1:** *Demographics for study population*. Study subjects are matched for age, sex, smoking (pack-yrs), caffeine (mg-yrs), NSAID use (mg-yrs), and head injuries (those with loss of consciousness or requiring medical attention). Chi-square was used for sex. Ordinary one-way ANOVA with Tukey’s adjustments for multiple comparisons was used for age, smoking, caffeine use, NSAID use, and head injuries. Welch’s t test was used for UPDRS motor score, Hoehn & Yahr scale, and disease duration. NHC neurologically healthy controls; RBD patients with REM sleep behavior disorder; EPD early-stage PD; MPD moderate-stage PD; NSAID nonsteroidal anti-inflammatory drug.

| **Table 1** | NHC | RBD | EPD | MPD | *p*-value |
| --- | --- | --- | --- | --- | --- |
| N | 21 | 15 | 27 | 31 |  |
| Age in years, mean (SD) | 65.29 (7.90) | 68.93 (4.90) | 67.19 (6.05) | 66.97 (6.27) | 0.4161 |
| Sex | 10M, 11F | 8M, 7F | 14M, 13F | 16M, 15F | 0.9865 |
| Smoking in pack-years, mean (SD) | 2.08 (6.01) | 7.05 (11.34) | 13.82 (21.24) | 9.00 (19.73) | 0.1298 |
| Caffeine in mg-years, mean (SD) | 9678 (6947) | 8124 (3869) | 8368 (4999) | 6636 (4745) | 0.2349 |
| NSAID use in mg-years, mean (SD) | 634.2 (1924) | 529.0 (1073) | 937.4 (1598) | 1187 (2326) | 0.6332 |
| Head injuries, mean (SD) | 0.81 (2.6) | 0.27 (0.46) | 0.48 (0.70) | 0.23 (0.50) | 0.4411 |
| UPDRS motor Score, mean (SD) | N/A | N/A | 14.08 (9.27) | 21.10 (10.72) | 0.0136 |
| Hoehn & Yahr scale, mean (SD) | N/A | N/A | 1.74 (0.44) | 2.04 (0.52) | 0.0494 |
| Disease duration from diagnosis in years, mean (SD) | N/A | N/A | 0.65 (0.75) | 5.69 (2.63) | <0.0001 |

To determine the effects of PD progression on stimulation-evoked innate immune responses, we first assessed the concentrations of inflammatory cytokines in the cultured media from isolated monocytes under baseline and stimulated conditions. Using multiplexed immunoassay platform (Meso Scale Discovery), we observed that IL-8 concentration in the media from vehicle-treated RBD monocytes was significantly reduced relative to other groups (Supplementary Fig. 1A), however absolute concentrations of TNF, IL-1β, and IL-10 were not significantly affected by PD status (Supplementary Fig. 1B-D). Due to significant variability in the data, we proceeded to normalize the concentrations of secreted cytokines in the stimulated condition to the amount secreted in the vehicle condition, allowing each patient to serve as their own normalization factor. We observed that monocytes from RBD patients exhibited significantly increased stimulation-dependent secretion of TNF, IL-1β, and IL-8 relative to all other groups (Fig. 2A-C). No significant differences were observed in IL-10 secretion between patient groups (Fig. 2D). Therefore, anti-inflammatory cytokine secretion was not as significantly modified by PD status as pro-inflammatory pathways. Collectively, such data support a pattern of upregulated stimulation-dependent secretion of proinflammatory cytokines by monocytes in the prodromal stage of PD, which subsequently diminishes as the disease progresses as evidenced by the levels in the early- and mid-stage PD cohorts.

**Fig. 2:**
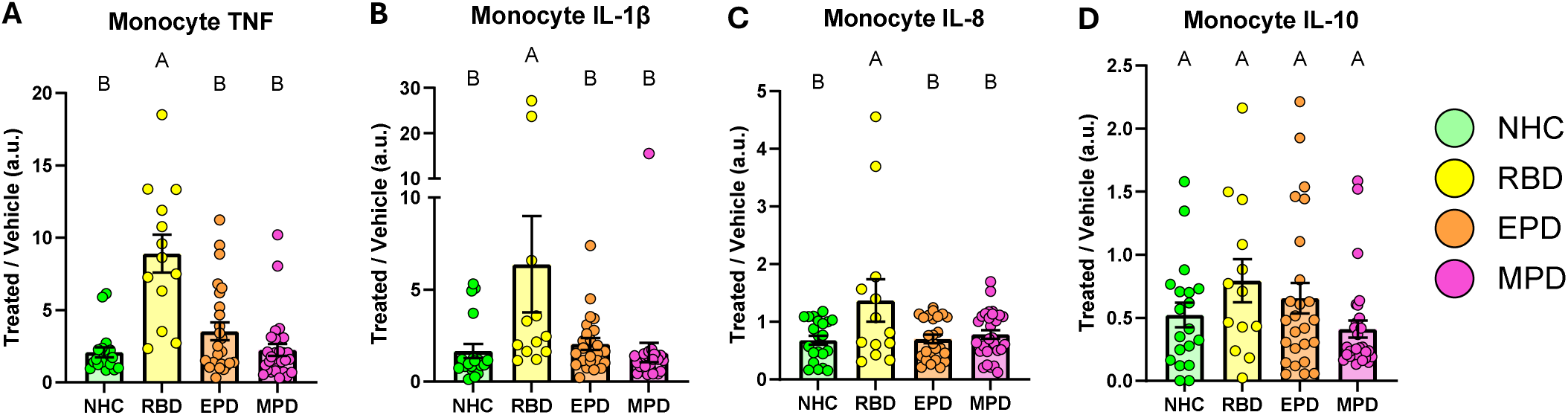
RBD patient monocytes exhibit increased stimulation-dependent cytokine secretion. Bar graphs overlaid with scatter plots showing stimulation-dependent secretion of inflammatory cytokines after IFNγ treatment from isolated monocytes from NHCs, RBD patients, EPD patients, and MPD patients. The stimulation-dependent secretion was quantified by normalizing the absolute concentration released in the stimulated condition to the absolute concentration released in vehicle for each participant. Stimulation-dependent secretion of TNF (**A**), IL-1β (**B**), IL-8 (**C**), and IL-10 (**D**). Bars represent mean +/- SEM. NHC neurologically healthy controls, *n* = 21; RBD patients with REM sleep behavior disorder, *n* = 15; EPD patients with early-stage PD, *n* = 27; MPD patients with moderate-stage PD, *n* = 30. Each symbol represents the measurement from a single individual. The results in **A-D** were analyzed using one-way ANOVA with Tukey’s corrections for multiple comparisons. Groups sharing the same letters are not significantly different (*p* > 0.05) whilst groups displaying different letters are significantly different (*p* < 0.05).

### Later stages of PD are characterized by progressive reductions in T cell cytokine secretion relative to individuals with RBD

We next sought to assess if isolated CD3^+^ T lymphocytes displayed similar or distinct patterns of stimulation-dependent cytokine secretion compared to those observed in monocytes. We began by comparing the absolute concentrations of cytokines in the cultured media of T lymphocytes following treatment with vehicle or CD3/CD28 Dynabeads. Absolute levels of TNF secreted from stimulated T cells from RBD patients were significantly increased relative to those from early and moderate PD (Supplementary Fig. 2A). This may indicate that the capacity for T lymphocytes to secrete TNF peaks during prodromal stages and diminishes upon the onset and progression of PD. In addition, we observed that IL-8 secretion from stimulated T cells was highest in early PD and reached statistical significance relative to NHCs (Supplementary Fig. 2B). Stimulated RBD T cells also showed increased secretion of IL-2 relative to moderate PD and increased secretion of IL-10 relative to NHC, but no differences were observed in IL-1β secretion across patient groups (Supplementary Fig. 2C-E). To mitigate variability, we again normalized the cytokine secretion from stimulated condition to the vehicle condition. Consistent with the analysis of absolute concentrations, relative stimulation-dependent secretion of TNF was increased from RBD T cells compared to early and moderate PD groups (Fig. 3A). No differences were observed across groups in terms of relative IL-1β and IL-8 secretion (Fig. 3B, C). Interestingly stimulation-dependent secretion of IL-2 from T cells was decreased in moderate PD relative to all other groups (Fig. 3D). In addition, stimulation-dependent IL-10 secretion was reduced in moderate PD relative to RBD patients (Fig. 3E). Overall, RBD T lymphocytes displayed a pattern of hyperinflammatory response to stimulation compared to moderate PD, which progressively diminishes as the stage of PD advances, in some cases dropping below NHC levels.

**Fig. 3:**
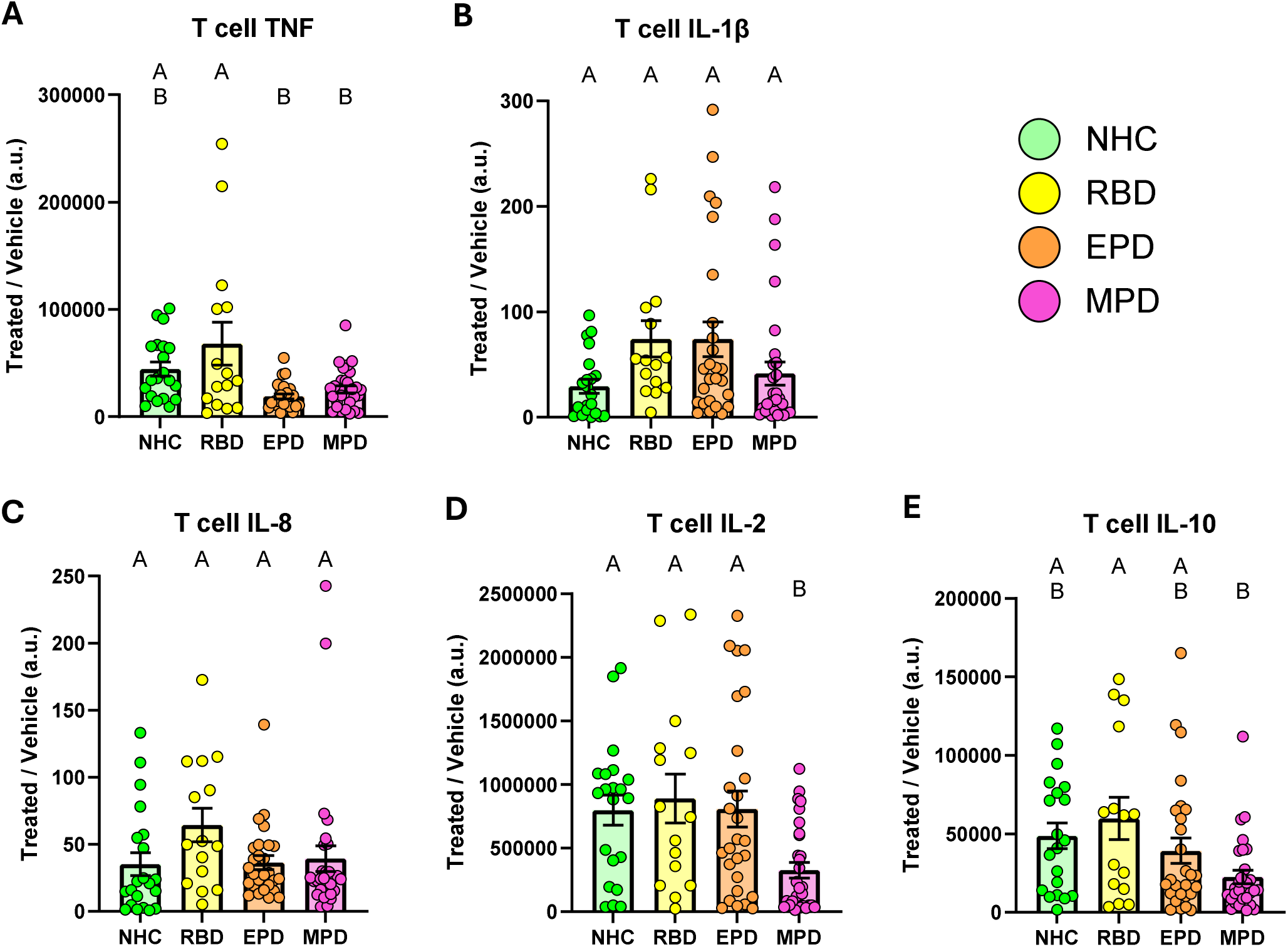
T cell stimulation-dependent cytokine secretion shows diminished response in moderate PD relative to RBD. Bar graphs overlaid with scatter plots showing stimulation-dependent secretion of inflammatory cytokines from isolated T lymphocytes after CD3/CD28 treatment from NHCs, RBD patients, EPD patients, and MPD patients. The stimulation-dependent secretion was quantified by normalizing the absolute concentration released in the stimulated condition to the absolute concentration released in vehicle for each participant. Stimulation-dependent secretion of TNF (**A**), IL-1β (**B**), IL-8 (**C**), IL-2 (**D**), and IL-10 (**E**). Bars represent mean +/- SEM. NHC neurologically healthy controls, *n* = 21; RBD patients with REM sleep behavior disorder, *n* = 15; EPD patients with early-stage PD, *n* = 27; MPD patients with moderate-stage PD, *n* = 30. Each symbol represents the measurement from a single individual. The results in **A-E** were analyzed using one-way ANOVA with Tukey’s corrections for multiple comparisons. Groups sharing the same letters are not significantly different (*p* > 0.05) whilst groups displaying different letters are significantly different (*p* < 0.05).

### Differences in PBMC subtype frequencies reveal changes in immunophenotype based on stage of PD

Prior studies have linked PD status to changes in the frequency of PBMC subtypes^8, 35^, so we sought to investigate if the observed changes in cytokine secretion were driven by changes in monocyte or T cell sub-population frequencies. To assess this, we stained the isolated monocytes and T cells after treatment for flow cytometry analysis using antibody-fluorophore conjugates for cell-surface markers (see Supplementary Figs. 3, 4 for gating strategy with fluorescence-minus-one controls (FMOCs)). We observed reduced frequencies of classical monocytes (CD14^+^CD16^-^) in vehicle-treated early and moderate PD groups relative to NHCs (Fig. 4A). IFNγ treatment caused an increase in the frequency of classical monocytes in all groups and ablated inter-group differences. Intriguingly, early PD patients displayed significantly elevated proportions of intermediate monocytes (CD14^+^CD16^+^) relative to NHCs in the vehicle condition (Fig. 4B). Furthermore, moderate PD patients showed higher frequencies of intermediate monocytes than both NHC and RBD groups in vehicle condition. This was surprising because our earlier data suggested that early and moderate PD monocytes as a whole display weaker stimulation-dependent cytokine secretion, yet intermediate monocytes are associated with strong secretion of proinflammatory cytokines^36^. We did not observe differences across patient groups in the frequency of non-classical (CD14^dim^CD16^+^) monocytes (Fig. 4C). IFNγ treatment evoked significant reductions in the frequencies of both intermediate and non-classical monocyte populations (Fig. 4B, C). These changes were not due to differences in the raw counts of total monocytes (Supplementary Fig. 5). In sum, these results suggest that PD is associated with a shift towards higher frequencies of intermediate monocytes at baseline, away from classical monocytes, but PD status does not interfere with the ability of monocytes to class switch between subtypes following immune stimulation.

**Fig. 4:**
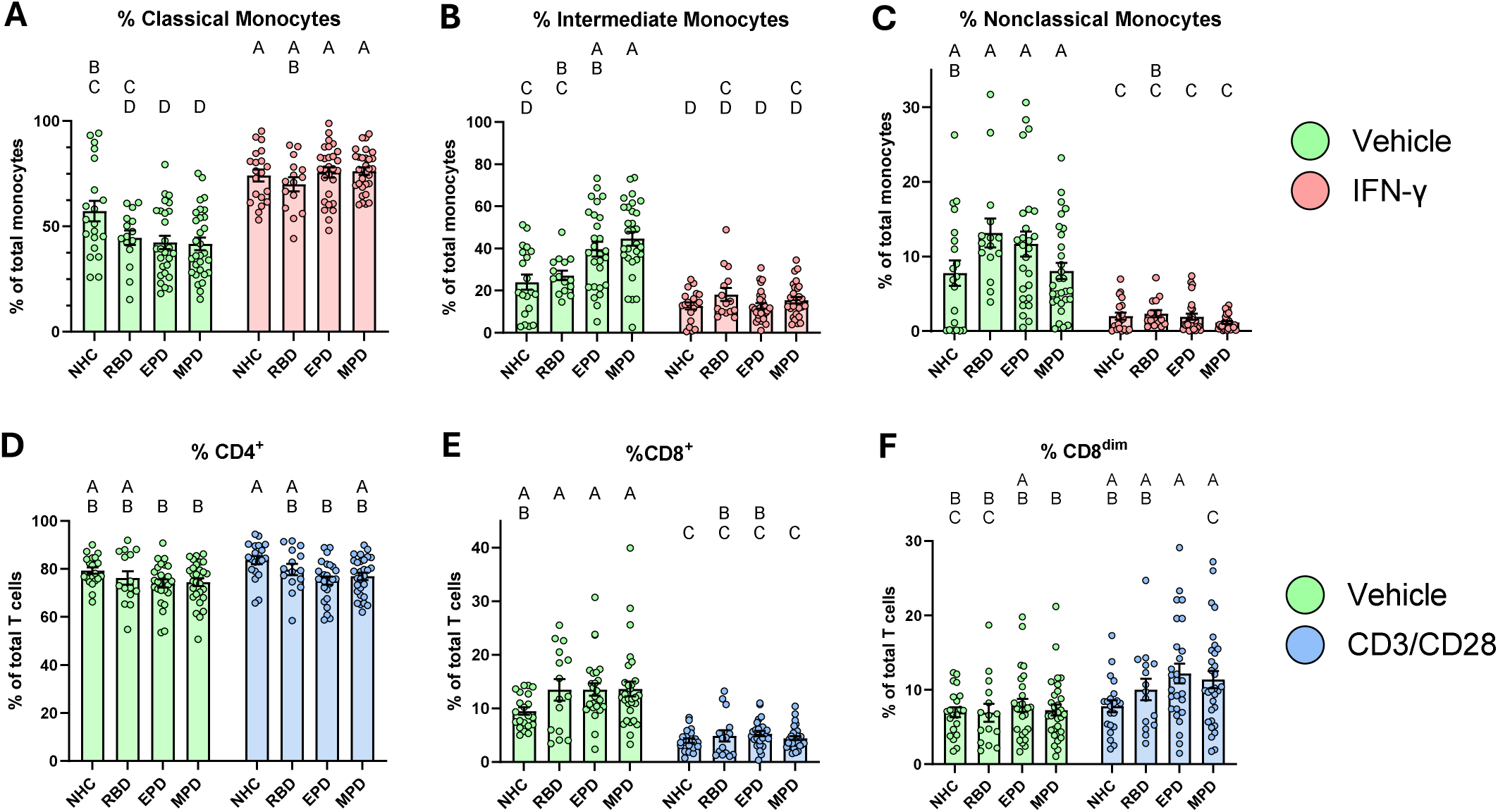
PBMC subtype frequencies across multiple stages of PD. Bar graphs overlaid with scatter plots showing the frequency of subtypes of monocytes and T cells in PBMCs from NHCs, RBD patients, EPD patients, and MPD patients. **A** Frequency of classical monocytes (CD14^+^CD16^-^) among total monocytes. **B** Frequency of intermediate monocytes (CD14^+^CD16^+^) among total monocytes. **C** Frequency of nonclassical monocytes (CD14^dim^CD16^+^) among total monocytes. **D** Frequency of CD4^+^CD8^-^ among total CD3^+^ T lymphocytes. **E** Frequency of CD4^-^CD8^+^ among total CD3^+^ T lymphocytes. **F** Frequency of CD4^-^CD8^dim^ among total CD3^+^ T lymphocytes. Bars represent mean +/- SEM. NHC neurologically healthy controls, *n* = 21; RBD patients with REM sleep behavior disorder, *n* = 15; EPD patients with early-stage PD, *n* = 27; MPD patients with moderate-stage PD, *n* = 30. Each symbol represents the measurement from a single individual. The results in **A-F** were analyzed using two-way ANOVA with Tukey’s corrections for multiple comparisons. Groups sharing the same letters are not significantly different (*p* > 0.05) whilst groups displaying different letters are significantly different (*p* < 0.05).

Next, we evaluated if the different stages of PD progression were associated with changes in the baseline or stimulation-evoked frequencies of CD4^+^ or CD8^+^ cells among T lymphocytes. CD4^+^ T cell frequencies were similar across groups in the vehicle treated condition, however CD3/CD28 stimulation resulted in a small but statistically significant decrease in CD4^+^ frequency in early PD relative to NHCs (Fig. 4D). The frequency of CD8^+^ T cells was not significantly different across groups in vehicle or stimulated conditions, and stimulation universally led to a decrease in the fraction of CD8^+^ cells (Fig. 4E). We also observed a population of CD4^-^ cells which were dimly positive for CD8, distinct from the brighter CD8^+^ population, and we termed this group CD8^dim^ (Supplementary Fig. 3). CD8^dim^ cells have been reported by others^37, 38^ and are believed to arise after excessive pathogen burden or immune activation^38^. Bead stimulation caused an increase in the frequency of CD8^dim^ T cells (treatment effect, *p* = 0.0001) (Fig. 4F), however, only the moderate PD group showed a statistically significant increase in CD8^dim^ after treatment. Downregulation of CD8 in T lymphocytes has been reported following exposure to viral and bacterial antigens^39^, and it has been hypothesized to promote peripheral tolerance^40, 41, 42^. Therefore, the increase in CD8^dim^ T cells after stimulation supports a pattern of increased peripheral tolerance in moderate-stage PD.

### Immune activation reveals deficits in CD8^+^ T cell mitochondrial health in moderate PD

Growing evidence suggests that cellular metabolic function is intricately connected to immune responses^43^. Given that decreased cytokine production in immune cells can be caused by mitochondrial dysfunction^44, 45^, we sought to assess if the stage of PD progression was associated with altered mitochondrial health in PBMC subsets. To investigate this, we quantified the median fluorescence intensity (MFI) of MitoTracker probes in stimulated PBMCs using flow cytometry. MitoTracker Green FM (MTG) was used to probe total mitochondrial content^46^. MitoTracker Red CMXRos (MTR) accumulates and stains mitochondria with sufficiently negative membrane potential^46^. A negative membrane potential is required for ATP production and normal mitochondrial function^47^, thus MTR MFI was used to quantify healthy mitochondrial content. We observed that total mitochondrial content and healthy mitochondrial content were not significantly different across patient groups in CD8^+^ T cells (Fig. 5A, B). To determine if PD progression modulates the relative proportion of healthy mitochondria within the cell, we calculated the MFI ratio of MTR/MTG and compared this ratio across groups. Intriguingly, stimulated CD8^+^ T cells from moderate PD patients showed significantly lower MTR/MTG ratio than NHCs (Fig. 5C). This effect was specific to stimulated cells, as no differences in MTR/MTG ratio were observed at baseline. In the CD8^dim^ population, we again did not observe an effect of PD status on total mitochondrial content or healthy mitochondrial content (Fig. 5D, E). However, CD8^dim^ lymphocytes from moderate PD patients showed significantly reduced relative mitochondrial health after stimulation (Fig. 5F), mirroring the findings from CD8^+^ cells. These results suggest that cytotoxic T cells from moderate PD patients are unable to maintain a high fraction of healthy mitochondria in response to immune activation and increased bioenergetic demand. We then examined CD4^+^ cells and observed that total mitochondrial content in RBD cells was significantly increased relative to all other groups in the vehicle condition (Fig. 5G). No differences were observed across groups in healthy mitochondrial content or relative mitochondrial health in CD4^+^ cells (Fig. 5H, I). These results suggest that RBD may be associated with a greater demand for mitochondria and energy production in CD4^+^ cells at baseline, however, RBD status does not disrupt the ability of these cells to regulate mitochondrial health in response to immune activation.

**Fig. 5:**
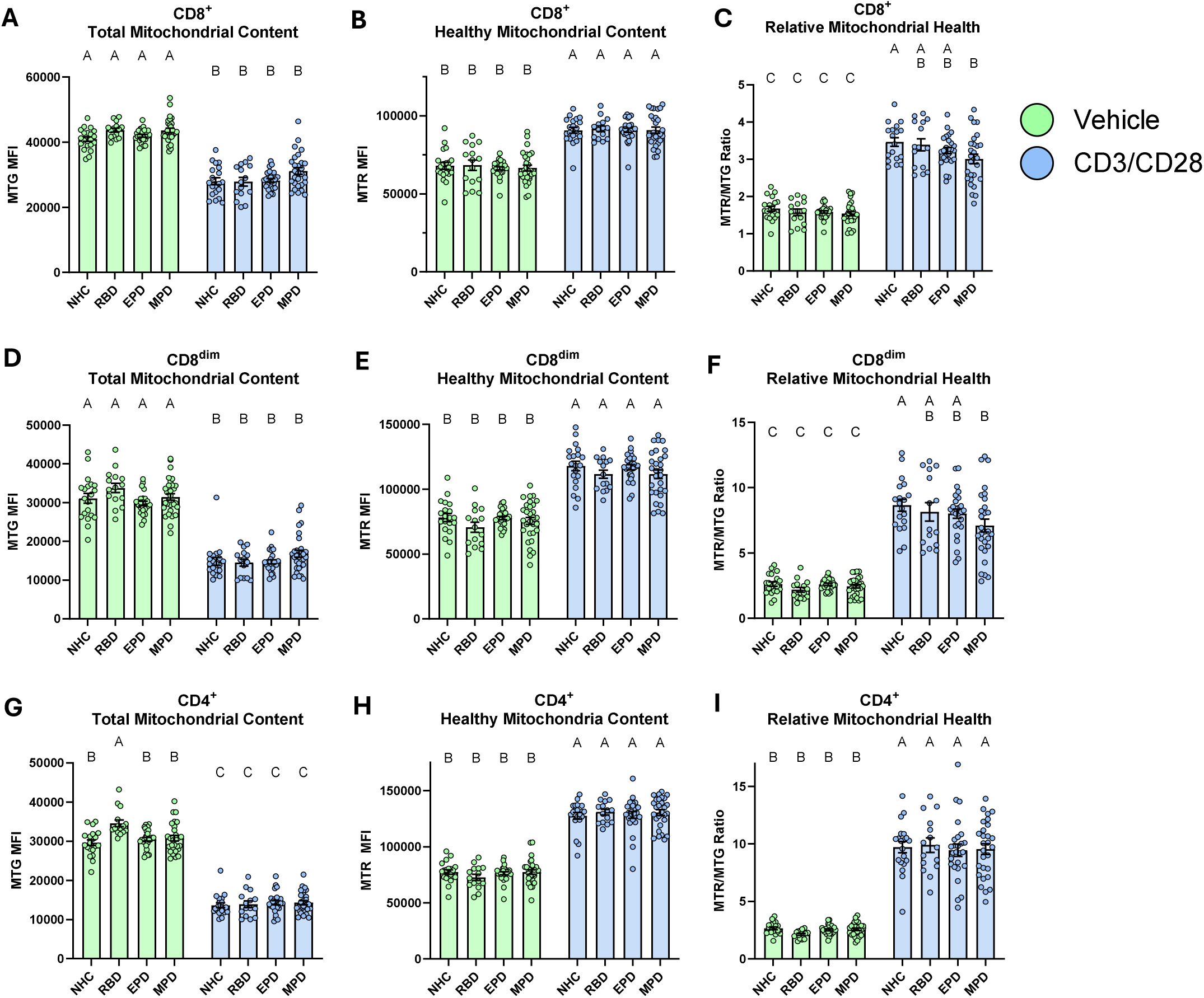
Cytotoxic T cells from moderate PD patients show impaired relative mitochondrial health following immune stimulation. Bar graphs overlaid with scatter plots showing the mitochondrial content and mitochondrial health after CD3/CD28 stimulation of T cell subsets from NHCs, RBD patients, EPD patients, and MPD patients. **A** Total mitochondrial content of CD8^+^ T lymphocytes. **B** Healthy mitochondrial content with negative membrane potential in CD8^+^ T lymphocytes. **C** Ratio of healthy mitochondrial content divided by the total in CD8^+^ T lymphocytes. **D** Total mitochondrial content of CD8^dim^ T lymphocytes. **E** Healthy mitochondrial content with negative membrane potential in CD8^dim^ T lymphocytes. **F** Ratio of healthy mitochondrial content divided by the total in CD8^dim^ T lymphocytes. **G** Total mitochondrial content of CD4^+^ T lymphocytes. **H** Healthy mitochondrial content with negative membrane potential in CD4^+^ T lymphocytes. **I** Ratio of healthy mitochondrial content divided by the total in CD4^+^ T lymphocytes. Bars represent mean +/- SEM. NHC neurologically healthy controls, *n* = 21; RBD patients with REM sleep behavior disorder, *n* = 15; EPD patients with early-stage PD, *n* = 27; MPD patients with moderate-stage PD, *n* = 30. Each symbol represents the measurement from a single individual. The results in **A-I** were analyzed using two-way ANOVA with Tukey’s corrections for multiple comparisons. Groups sharing the same letters are not significantly different (*p* > 0.05) whilst groups displaying different letters are significantly different (*p* < 0.05). MTG MitoTracker Green FM, MTR MitoTracker Red CMXRos.

Next, we examined mitochondrial health in monocyte subpopulations, and we observed that PD status was not associated with significant changes in mitochondrial health in classical and intermediate monocytes (Supplementary Fig. 6A-F). In nonclassical monocytes, total mitochondrial content and healthy mitochondrial content also remained unaffected by PD status (Supplementary Fig. 6G, H), but the relative mitochondrial health in moderate PD patients was significantly reduced compared to early PD after stimulation. These data suggest that PD status does not compromise the ability of monocytes to regulate mitochondrial health after stimulation, with the exception of nonclassical monocytes in late stages of disease.

### Relative mitochondrial health in CD8^+^ T cells is positively correlated with stimulation-dependent T cell cytokine secretion across the PD spectrum

Previous reports show that mitochondrial deficits in immune cells can contribute to decreased cytokine secretion^44, 48^. Therefore, we sought to explore if the deficits we observed in mitochondrial health subsequent to immune stimulation were correlated with deficient cytokine secretion across multiple stages of PD. We had noted that reductions in relative mitochondrial health with advanced stages of PD were specific to cytotoxic T cells, thus we used linear regression to determine the relationship between CD8^+^ MTR/MTG ratio and T cell cytokine secretion. Relative mitochondrial health was found to be significantly positively correlated with T cell stimulation-dependent secretion of TNF, IL-8, IL-2, and IL-10 (Fig. 6A-D) but not IL-1β (Fig. 6E). To ensure this relationship was not driven by motor severity, we also compared patient UPDRS scores in early and moderate PD groups to cytokine secretion and observed no statistically significant correlations (Supplementary Fig. 7). These results indicate that mitochondrial health may represent a useful metric for understanding peripheral immune responses across multiple stages of PD progression, including in RBD patients.

**Fig. 6:**
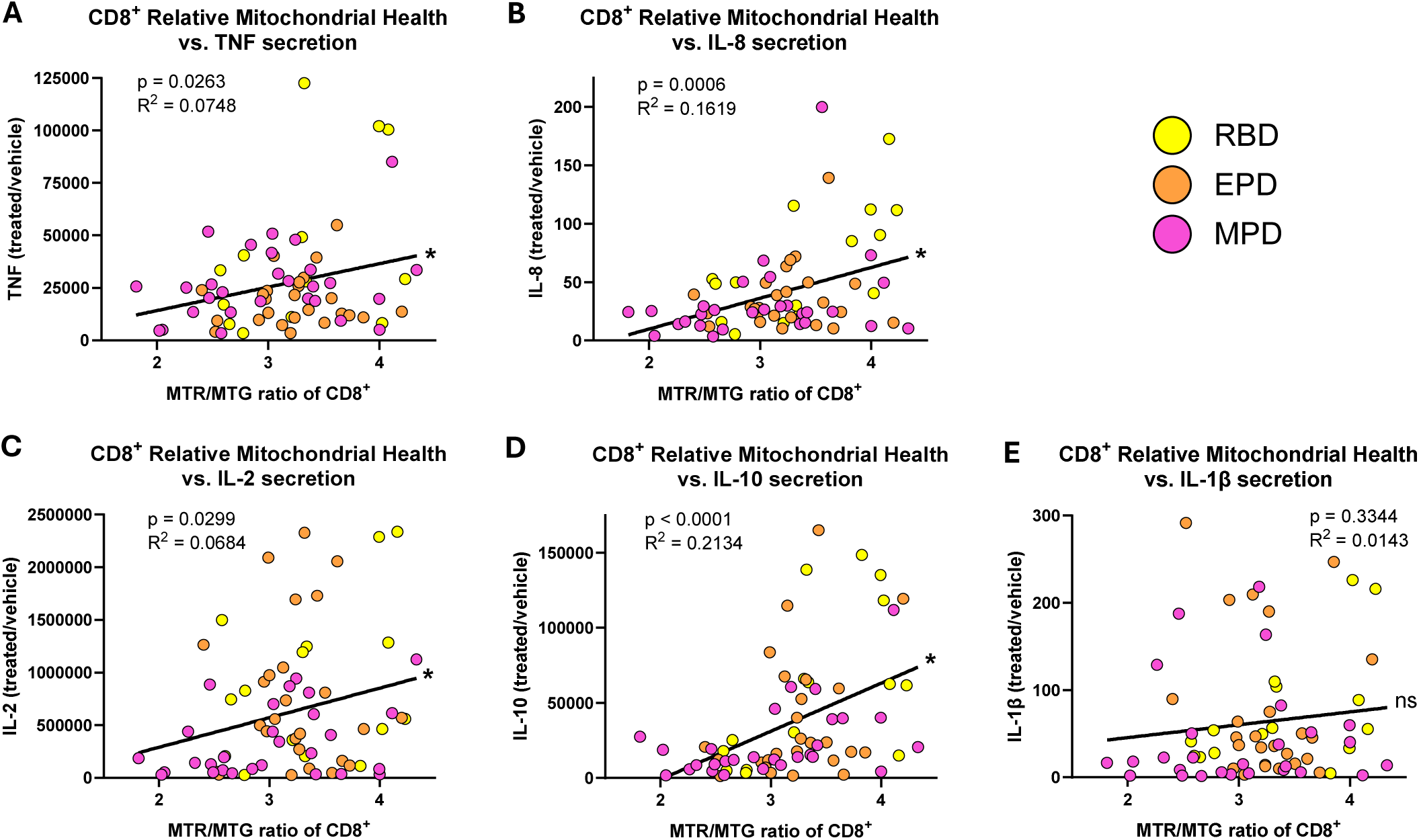
Relative mitochondrial health in CD8+ T cells is positively correlated with stimulation-dependent cytokine secretion from T cells in disease. Correlational analysis showing the relationship between relative mitochondrial health of CD8^+^ T cells and stimulation-dependent cytokine secretion from total T cells in RBD patients, EPD patients, and MPD patients. Correlation between relative mitochondrial health after CD3/CD28 Dynabead stimulation and stimulation-dependent secretion of TNF (**A**), IL-8 (**B**), IL-2 (**C**), IL-10 (**D**), and IL-1β (**E**). Coefficient and *p*-value based on Pearson correlation. Each symbol represents the measurement from a single individual. RBD patients with REM sleep behavior disorder, *n* = 15; EPD patients with early-stage PD, *n* = 27; MPD patients with moderate-stage PD, *n* = 30. * signifies that the slope of the line is significantly different from zero (*p* < 0.05).

### Monocyte lysosomal function and LRRK2 activity differentiate RBD from motor PD

PD has been linked to deficits in lysosomal degradation in PBMCs^49^, and the severity of motor symptoms in PD patients is inversely correlated with monocyte activity of the lysosomal enzyme glucocerebrosidase^33^. Furthermore, pharmacological inhibition of lysosomal function polarizes immune cells towards a more proinflammatory phenotype and enhances proinflammatory cytokine production^50^. This led us to question whether different stages of PD were associated with stimulation-dependent changes in lysosomal health in PBMC subsets. To explore this, the MFI of Lysotracker Red DND-99 (LTR), which accumulates in and stains sufficiently acidic lysosomes within cells, was quantified in monocytes and T cells after treatment with our stimulation paradigm. We observed that lysosomal content, quantified from LTR MFI, was not significantly affected by PD status in monocytes, CD4^+^ T cells, or CD8^+^ T cells (Supplementary Fig. 8A-E). However, CD8^dim^ T cells from RBD patients showed increased lysosomal content relative to all other groups in vehicle treated condition (Supplementary Fig. 8F). Next, we used the pan-cathepsin fluorescent probe BMV109 to quantify lysosomal cathepsin activity as a marker of lysosomal function in these cells. We first observed that classical monocytes from RBD patients exhibited increased cathepsin activity relative to NHCs in vehicle condition (Fig. 7A). Additionally, intermediate monocytes from RBD patients showed increased cathepsin activity in vehicle condition relative to NHCs (Fig. 7B). No differences in cathepsin activity were observed across groups in nonclassical monocytes (Fig. 7C) or T cell populations (Supplementary Fig. 9). Collectively, these data suggest that PD is associated with an upregulation of baseline lysosomal degradative capacity in specific monocyte subsets, and this upregulation can be detected in the prodromal stage of disease.

**Fig. 7:**
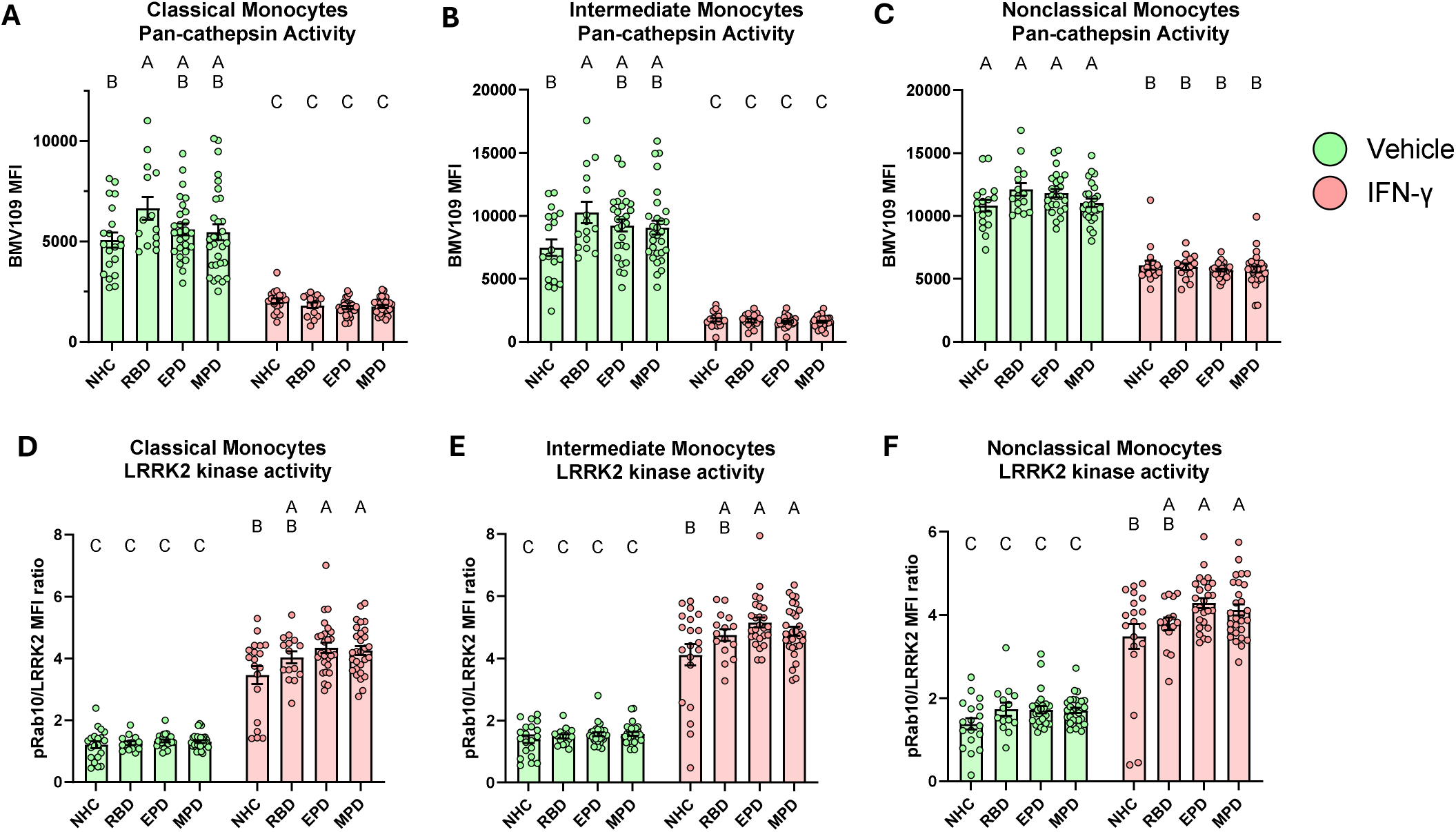
Monocyte subpopulations display distinct profiles of pan-cathepsin and LRRK2 kinase activity based on PD stage. Bar graphs overlaid with scatter plots showing the lysosomal pan-cathepsin activity and LRRK2 kinase activity in monocyte subtypes after IFNγ treatment from NHCs, RBD patients, EPD patients, and MPD patients. **A** Pan-cathepsin activity of classical monocytes (CD14^+^CD16^-^). **B** Pan-cathepsin activity of intermediate monocytes (CD14^+^CD16^+^). **C** Pan-cathepsin activity of nonclassical monocytes (CD14^dim^CD16^+^). **D** pRab10 expression normalized to LRRK2 expression in classical monocytes (CD14^+^CD16^-^). **E** pRab10 expression normalized to LRRK2 expression in intermediate monocytes (CD14^+^CD16^+^). **F** pRab10 expression normalized to LRRK2 expression in nonclassical monocytes (CD14^dim^CD16^+^). Bars represent mean +/- SEM. NHC neurologically healthy controls, *n* = 21; RBD patients with REM sleep behavior disorder, *n* = 15; EPD patients with early-stage PD, *n* = 27; MPD patients with moderate-stage PD, *n* = 30. Each symbol represents the measurement from a single individual. The results in **A-F** were analyzed using two-way ANOVA with Tukey’s corrections for multiple comparisons. Groups sharing the same letters are not significantly different (*p* > 0.05) whilst groups displaying different letters are significantly different (*p* < 0.05). pRab10 phosphorylated Rab10, LRRK2 leucine-rich repeat kinase 2.

Alterations in leucine-rich repeat kinase 2 (*LRRK2*) expression have been heavily implicated in lysosomal biology and function^51, 52^, and the gain-of-kinase mutation *LRRK2-*G2019S is one of the most common causes of genetic PD^53^. Additionally, our previous work described changes in the expression of both LRRK2 and phosphorylated Rab10 (pRab10), a LRRK2 substrate indicative of kinase activity, in peripheral immune cells from iPD patients^34^. Recent observations have also defined a role for LRRK2 in suppressing lysosomal degradative activity and regulating cytokine secretion^34, 54^. In light of this evidence, we sought to explore the effects of PD progression on LRRK2 expression and kinase activity in stimulated PBMC subsets. We found that IFNγ treatment increased both LRRK2 and pRab10 expression in all monocyte subtypes (Supplementary Fig. 10). Relative LRRK2 kinase activity was assessed by normalizing pRab10 MFI to LRRK2 MFI, and this revealed significantly increased stimulation-dependent LRRK2 kinase activity in all monocyte subtypes from early PD and moderate PD groups relative to NHCs (Fig. 7D-F). Meanwhile, LRRK2 expression and kinase activity in T cell subsets were similar across different stages of PD (Supplementary Fig. 11). Jointly, these experiments indicate that PD is associated with increases in stimulation-dependent LRRK2 activity, and these effects are cell-type specific.

## Discussion

Emerging literature suggests that immunophenotypic assays have the potential to more clearly define the extent of peripheral immune dysregulation in PD, thereby revealing novel biomarkers and tools for early diagnosis. However, the existence of dysregulated cytokine secretion prior to the onset of motor symptoms in PD remains a matter of contention^19, 20^. Furthermore, studies thus far have overlooked the potential for cell-type specific differences in cytokine secretion, leaving open the possibility that specific PBMC subsets exhibit distinct patterns of immune dysfunction in PD. Here we showed that isolated monocytes and T cells from RBD patients display a distinct pattern of immune dysregulation, and that immunometabolic responses to stimulation stratify multiple stages of PD progression. Our methodology revealed cell-type specific differences in stimulation-evoked cytokine secretion, PBMC population composition, and facets of mitochondrial and lysosomal health which change dynamically with disease stage.

Our work provides the first evidence that stimulation-evoked cytokine secretion from RBD monocytes is increased relative to NHC and PD groups. Previous studies have investigated circulating cytokine levels in RBD patients and reported heterogenous results^19, 20^, however these studies did not examine the *ex vivo* response to stimulation nor did they include early or moderate iPD patients. Our analysis suggests that baseline levels of cytokine secretion are not significantly altered in RBD monocytes, but instead that these patients display an aberrant upregulation of TNF, IL-1β, and IL-8 secretion in response to immune stimuli. This may lead to increased peripheral inflammation and represent a potential mechanism which contributes to PD conversion and progression. Support for this comes from epidemiological reports that anti-TNF therapy in inflammatory bowel disease patients and ibuprofen use in the general population are associated with lower incidence of PD in these groups^55, 56^. Ongoing clinical trials for RBD and PD include biofluid measurements to assess inflammation^57, 58^, however, we propose that stimulation-evoked cytokine secretion may have greater sensitivity and will more accurately report on the cell-type specificity and efficacy of immunomodulatory treatments. Thus, the inclusion of stimulation-dependent immune responses *ex vivo* will be necessary in future studies to directly investigate peripheral immune dysregulation as a contributing mechanism in PD pathogenesis and progression.

Concurrently, we observed that T lymphocytes from moderate PD patients exhibit reduced secretion of TNF, IL-2, and IL-10 relative to RBD patients, a pattern suggestive of relative immune exhaustion^59^. Given that IL-2 controls regulatory T cell maturation and proliferation^60^, our results suggest that regulatory T cell maturation may be disrupted in moderate PD consistent with current literature^61^. We also show that cytotoxic T cells in moderate PD display poorer relative mitochondrial health after stimulation compared to NHCs, and furthermore, that relative mitochondrial health after stimulation is significantly correlated with cytokine secretion. Mitochondrial impairment in neurons is widely reported in idiopathic and genetic forms of PD^62, 63, 64^. Given that reduced mitochondrial respiration is sufficient to elicit T cell exhaustion in chronic infections^44^, this may explain the immune exhaustion-like phenotype observed in moderate PD cells. Notably, UPDRS did not correlate with stimulation-dependent cytokine secretion, suggesting that motor severity is not sufficient to capture the underlying biology of PD as the disease progresses. Current perspectives acknowledge that additional metrics beyond motor progression are necessary to appreciate emerging sub-types of PD^65, 66, 67^, and a combination of biochemical markers with clinical measures may be necessary to precisely identify patient endophenotypes for reduced heterogeneity in clinical trials^68, 69^.

Our analysis revealed a hyper-inflammatory response in peripheral immune cells from RBD patients and immune exhaustion in later PD stages. Immunogenic mitochondrial damage-associated molecular patterns (DAMPs) and mitochondrial DNA are known to activate TLR4^70, 71^, and these could be present at higher concentrations in peripheral immune cells from RBD patients. In support of this, a recent study found that fibroblasts from RBD patients who converted to PD had significantly increased mitochondrial fragmentation relative to controls, and isolated RBD patients showed similar but milder alterations^72^. Therefore, we posit that mitochondrial fragmentation predisposes towards a hyperinflammatory response early in disease course, but persistent accumulation of mitochondrial dysfunction eventually overwhelms the adaptive ability of these cells and leads to immune exhaustion later in PD. Future research should seek to evaluate mitochondrial morphology in RBD and PD immune cells, as pharmacologic interventions aimed at improving mitochondrial biogenesis and enhancing mitophagy represent novel and exciting avenues that should be explored to delay disease progression.

Flow cytometry analysis revealed that moderate PD patients show a reduced frequency of classical monocytes and increased frequency of intermediate monocytes compared to NHCs. Our findings are consistent with those from Thome et al. who reported identical changes in monocyte subtype frequencies in PD patients^61^. Intermediate monocytes have been reported to secrete higher levels of proinflammatory cytokines than classical monocytes in some contexts^73, 74^, which seems at odds with our findings that showed moderate PD monocytes have diminished stimulation-dependent cytokine secretion relative to RBD. One potential explanation is that the shift towards increased numbers of intermediate monocytes is a compensatory response to underlying functional deficits in these individuals. In addition, we observed that immune stimulation of T lymphocytes led to an increased frequency of CD8^dim^ cells in most groups; however this only reached statistical significance in moderate PD patients. Downregulation of CD8 in T lymphocytes has been reported following exposure to viral and bacterial antigens^39^ and has been hypothesized to promote peripheral tolerance^40, 41, 42^. Consequently, moderate PD may be associated with increased peripheral tolerance in cytotoxic T cells after immune stimulation. Our data so far cannot differentiate more specialized cell types such as helper, regulatory, or natural killer T cells, which will be important to comprehensively describe how PD stage affects peripheral immune populations. Thus, further efforts using scRNAseq or similar methodologies capable of garnering deep cell-type specific data to explore differences across these PBMC subsets should be considered.

Our study also demonstrated increased pan-cathepsin activity in classical and intermediate monocytes from RBD patients at baseline relative to NHCs. Upregulation of cathepsins has been reported in multiple PD models^75, 76^; however our work provides the first evidence that lysosomal function as measured by pan-cathepsin activity is upregulated in RBD patients. Cathepsin activity is known to modulate autophagic flux^77^, therefore this may represent an increased demand for lysosomal degradation and autophagy in the prodromal stage of PD. Indeed, autophagic flux increases to compensate for mitochondrial defects^78^, and mitochondrial membrane potential is a known regulator of autophagic flux^79^. Lysosomal and mitochondrial dynamics are extensively linked to one another, so additional research is necessary to determine if poor mitochondrial health in RBD is a downstream consequence of disturbances in lysosomal activity. In sum, our findings point to observable changes in metabolic organelle function in RBD patients, suggesting that peripheral immunometabolic deficits are present in the prodromal stages of PD. Moreover, these deficits could be ascertained non-invasively to stratify and enroll patients who are more likely to benefit from interventions targeting cellular metabolism.

In addition, we observed a mild increase in stimulation-dependent LRRK2 kinase activity from RBD monocytes relative to NHCs, although this did not reach statistical significance. A larger and significant increase was observed with early and moderate PD, creating an overarching pattern consisting of gradual increases in LRRK2 kinase activity as the diseases progresses. While inhibition of LRRK2 kinase activity has been proposed as a potential approach to combat neuroinflammation in iPD^80^, our data would suggest that this may be of limited benefit in modulating peripheral inflammation in RBD patients. Instead, inhibiting LRRK2 kinase activity may be more beneficial specifically for individuals carrying the G2019S gain-of-kinase mutation, because we have previously demonstrated that LRRK2-targeting anti-sense oligonucleotide therapy reduces inflammatory cytokine secretion in *LRRK2-*G2019S peripheral macrophages^81^. However, our results cannot conclusively distinguish whether increased LRRK2 kinase activity is compensating for or contributing to immune dysregulation in the prodromal stage of iPD, therefore additional *ex vivo* experiments with pharmacologic inhibitors are warranted.

Our work reveals immunophenotypic differences which could serve as the basis for stratification of RBD patients with prodromal PD. Individuals with prodromal PD whose signature of immune activation is distinct from patients with motor manifestations of PD would likely benefit from early immunomodulatory interventions to arrest or slow progression. However, some limitations to this strategy must be noted. The estimated risk for patients with isolated RBD to develop PD is 43%, with the next most likely course being a 25% risk for dementia with Lewy bodies^2^. Therefore, RBD is not completely specific for prodromal PD, and our findings may represent dysregulation in a common pathway shared by multiple neurodegenerative diseases. Second, our study design is based on cross-sectional analyses and did not include longitudinal follow up or sampling for the enrolled RBD patients; thus, we cannot conclusively determine which of these patients will ultimately convert to PD. It will be vital for future longitudinal studies to be conducted with sufficient power to replicate these findings and correlate these immune markers with eventual PD conversion rates. In addition, our findings suggest that T cells from moderate PD patients exhibit deficits in production of cytokines such as IL-2 which is generally consistent with an immune exhaustion phenotype^59^. However, further evaluations are required to definitively make the distinction between immune exhaustion and senescence. In line with this, evaluations of cell-surface markers of exhaustion such as PD-1, CTLA-4, and TIM-3 are underway in parallel studies.

In summary, this work demonstrates that peripheral blood immune cells from RBD patients display a distinct pattern of stimulation-dependent immune dysregulation relative to NHCs and clinically diagnosed PD groups. These findings hold significant potential to advance both scientific understanding and clinical practice for PD. First, our use of sensitive stimulation-dependent assays and focus on cell-type specific cytokine secretion has allowed us to detect a specific pattern of peripheral immune dysregulation in RBD patients, thereby addressing inconsistencies in previous studies. Second, our finding that T cell mitochondrial health correlates with stimulation-dependent cytokine secretion across the disease spectrum reveals a novel target for therapeutic intervention. Early intervention to rescue mitochondrial deficits may help mitigate excessive inflammation in RBD patients, while in advanced stages it may be a crucial target to combat immune exhaustion or senescence. Lastly, we recommend the expanded use of *ex vivo* stimulation based peripheral immune cell biomarkers, such as measures of mitochondrial health, to distinguish unique immunophenotypes for PD patients. It is becoming increasingly apparent that disease duration and severity of motor symptoms are not sufficient to describe the spectrum of PD progression, and the inability of the field to capture this heterogeneity across patients has been a barrier to success for clinical trials. The insights from our data should open up promising avenues for future research as the field continues to search for clinically relevant biomarkers. Moreover, applying these findings to clinical practice has the potential to significantly enhance patient-centered precision medicine in iPD.

## Materials and Methods

### Human Subjects

This study was reviewed and approved by the University of Florida Institutional Review Board (IRB202002639). Participants provided written informed consent to participate. Blood was initially collected from healthy volunteers to establish and optimize assay parameters. Then 15 subjects with RBD, 27 subjects with early PD, 30 subjects with moderate PD, and 21 age-matched, neurologically normal control subjects were recruited through the Norman Fixel Institute for Neurological Diseases at the University of Florida for this study. Early PD patients were <2 years post-diagnosis with <1 year on PD medications. Moderate PD patients were between 2-10 years post-diagnosis. Subjects were excluded based on age (younger than 30 and over 80 years of age), known familial PD mutations and/or other known neurological, chronic or recent infections, or autoimmune comorbidities. Subjects were genotyped for the *LRRK2*-G2019S mutation (Life Technologies #4351378, Grand Island, NY) and excluded from this study if they were shown to be mutation carriers.

During recruitment, a family history and environmental questionnaire was used to assess history of disease and inflammation/immune-relevant environmental exposures and comorbidities. Caffeine use, non-steroidal anti-inflammatory drug (NSAID) use, and nicotine exposure was calculated as mg-years, mg-years, and pack-years, respectively. The study populations were balanced with respect to risk factors for PD, including age, smoking, non-steroidal anti-inflammatory drug use, and caffeine intake (Table 1).

### Peripheral Blood Mononuclear Cell (PBMC) Isolation and Cryopreservation

Cell isolation was accomplished using BD Vacutainer CPT Cell Preparation Tube with Sodium Citrate (BD Biosciences, 362761). Approximately 6 CPT tubes, each containing 8 mL of blood, were collected from each participant. CPT tubes were inverted 8–10 times and centrifuged at room temperature at 1500 x *g* for 20 min at room temperature. The PBMC enriched layer was transferred to a new 50 mL conical tube and MACS buffer (PBS, 0.5% bovine serum albumin, 20 mM EDTA, pH 7.2) was added to a final volume of 50 mL, followed by centrifugation at 1800 x *g* for 10 min at room temperature. Following removal of the supernatant, PBMCs were resuspended in 10 mL MACS buffer and counted on a hemocytometer using Trypan blue (1:20 dilution) exclusion to ascertain viability.

Next, to cryopreserve the samples, PBMCs were centrifuged for 5 min 1800 x *g* at room temperature. Supernatant was aspirated and cell pellets were gently resuspended in cryopreservation media (54% RPMI 1640, 36% FBS, 10% DMSO) at a final concentration of 1 × 10^7^ cells/mL in cryovials (Simport, T311-2). Cryovials were placed in a room-temperature Mr. Frosty freezing container with isopropanol as per manufacturer’s instructions and stored at −80°C overnight. After overnight storage at −80°C, the next day cryovials were removed from freezing containers and placed into liquid nitrogen for long-term storage.

### Cryorecovery of Isolated PBMCs

For cryorecovery, cryovials of PBMCs were retrieved from liquid nitrogen, rapidly thawed in a water bath at 37°C, and rapidly added to 25 mL of 37°C filter sterilized complete culture media (RPMI 1640 media, 10% low endotoxin heat-inactivated FBS, 1 mM Penicillin-Streptomycin). PBMCs were pelleted via centrifugation at 300 x *g* for 10 min at room temperature. Pellets were gently resuspended in 10 mL of 37°C MACS buffer (PBS, 0.5% bovine serum albumin, 20 mM EDTA, pH 7.2), then viability and cell count were obtained with a hemocytometer using Trypan blue (1:20 dilution) exclusion to ascertain viability.

### *Ex vivo* Isolation of CD3+ T cells and Pan Monocytes from PBMCs

Following cryorecovery, CD3^+^ T cells were isolated from total PBMCs using REAlease® CD3 MicroBead Kit, human (Miltenyi, 130-117-038) following the manufacturer’s instructions with slight modifications. PBMCs were centrifuged at 300 x *g* for 10 min at room temperature, supernatant was aspirated, and pellets were gently resuspended in 40 μL of separation buffer (PBS, 0.5% bovine serum albumin, 2mM EDTA, pH 7.2) per 1x10^7^ total cells. 10 μL of REAlease CD3-Biotin were added per 1x10^7^ total cells, mixed well, and samples were incubated at room temperature for 5 minutes. 100 μL of REAlease Anti-Biotin Microbeads (CD3, human) were added per 1x10^7^ total cells, mixed well, and samples were incubated at room temperature for 5 minutes. Samples were diluted to a total volume of 2 mL with separation buffer then passed through pre-wetted LS columns (Miltenyi, 130-042-401) in a QuadroMACS™ Separator (Miltenyi, 130-091-051). Columns were washed 3 times with 3 mL of separation buffer, and the flow-through was set aside at 4°C for isolation of monocytes. LS columns were removed from the magnetic separator and flushed twice with 5 mL of REAlease Bead Release buffer to release bead-bound CD3^+^ cells. CD3^+^ samples were mixed well and incubated at room temperature for 5 minutes. Then, CD3^+^ samples were centrifuged at 300 x *g* for 10 min at 4°C, supernatant was aspirated, pellets were gently resuspended in 5 mL separation buffer. 100 μL of REAlease Release Reagent was added to each sample, mixed well, and then CD3^+^ cells were counted using a hemocytometer with Trypan blue (1:20 dilution) to ascertain viability.

Monocytes were isolated from the flow-through of CD3^-^ cells using Pan Monocyte Isolation Kit, human (Miltenyi, 130-096-537) following the manufacturer’s instructions with slight modifications. Cells were centrifuged at 300 x *g* for 10 min at 4°C, supernatant was aspirated, and pellets were gently resuspended in 45 μL of cold separation buffer per 1x10^7^ total cells. 15 μL of FcR blocking reagent and 18.75 μL of Biotin-antibody cocktail was added per 1x10^7^ total cells, samples were mixed well, and then cells were incubated for 5 minutes at 4°C. 45 μL of cold separation buffer and 30 μL of Anti-Biotin Microbeads were added per 1x10^7^ total cells, samples were mixed well, and then cells were incubated for 5 minutes at 4°C. Samples were diluted to a total volume of 2 mL with cold separation buffer then passed through pre-wetted LS columns in a QuadroMACS™ Separator. Columns were washed 3 times with 3 mL of separation buffer, and the flow-through containing purified monocytes was counted on a hemocytometer using Trypan blue (1:20 dilution) to ascertain viability.

### *Ex vivo* T Cell and Monocyte Cell Culture Plating and Treatments

T cells were diluted to a final concentration of 1x10^6^ per mL in 1mL complete culture media in 24-well plates and allowed to rest for 2 hours at 37°C, 5% CO2, 95% relative humidity. After resting, cells were treated with either vehicle or 3.125 μL Dynabeads™ Human T-Activator CD3/CD28 (Gibco, 11161D) for 72 hours at 37°C, 5% CO2, 95% relative humidity.

Monocytes were diluted to a final concentration of 5x10^5^ per mL in 1mL complete culture media in 24-well plates and allowed to rest for 2 hours at 37°C, 5% CO2, 95% relative humidity. After resting, cells were treated with either vehicle or 200U human IFNγ (Peprotech, 300-02) for 72 hours at 37°C, 5% CO2, 95% relative humidity.

### Live Cell Flow Cytometry Assay for Mitochondrial Health and Pan-cathepsin Activity

After the 72-hour stimulation, cells were harvested and centrifuged at 300 x *g* for 10 min at 4°C. Supernatant was collected to quantify cytokine secretion (described below). Cell pellets were gently resuspended in 200 μL of cold PBS and transferred to a v-bottom 96-well plate (Sigma, CLS3896-48EA). Samples were centrifuged at 300 x *g* for 5 min at 4°C. Cells were resuspended in 200 μL of complete growth media containing 1 μM MitoTracker™ Red CMXRos (Invitrogen, M7512),1 μM MitoTracker™ Green FM (Invitrogen, M7514), and 1 μM BMV109 Pan Cathepsin Probe (Vergent Biosciences, 40200-200). Cells were incubated for 1 hour at 37°C in the dark. Samples were centrifuged at 300 x *g* for 5 min at 4°C. Cell pellets were resuspended in PBS and washed x 2 by centrifugation at 300 x *g* for 5 min at 4°C. Cells were resuspended in 50 μL of Live/Dead Fixable Violet stain (diluted 1:2000 in PBS, Invitrogen, L34962) and incubated in the dark at room temperature for 30 min. Cells were centrifuged at 300 x *g* for 5 min at 4°C washed in PBS x 2. Cells were resuspended in 50 μL of PBS containing diluted antibodies (see **Table 2** for T cell panel, see **Table 3** for monocyte panel) and incubated in the dark at 4°C for 20 min. Cells were centrifuged at 300 x *g* for 5 min at 4°C washed in FACS buffer (PBS, 0.5 mM EDTA, 0.1% sodium azide) x 3. Cells were analyzed via flow cytometry on a FACSymphony™ A3 cytometer (BD Biosciences). Data were analyzed using FlowJo version 10.10.0 software (BD Biosciences). When validating all flow cytometry panels and antibodies, fluorescence minus one controls (FMOCs) were used to set gates and isotype controls were used to ensure antibody-specific binding.

**Table 2:** Flow cytometry T cell panel marker antibody panel. This antibody panel was used for both unfixed and fixed cell panels to identify cell-surface markers of T cell subsets.

| **Table 2** |  |  |  |  |
| --- | --- | --- | --- | --- |
| Flow-cytometry T cell marker antibody panel |  |  |  |  |
| **Target** | **Conjugate** | **Catalog #** | **Dilution** | **Company/Manufacturer** |
| CD3 | BUV737 | BDB612752 | 1:50 | BD Biosciences |
| CD4 | BUV395 | BDB564724 | 1:50 | BD Biosciences |
| CD8 | BV605 | BDB564116 | 1:50 | BD Biosciences |
| CD137 | Pe-Cy7 | 309818 | 1:20 | BioLegend |
| FcR block | - | 422301 | 1:20 | BioLegend |
| Live/dead Violet | - | L34962 | 1:2000 | Invitrogen |

**Table 3:** Flow cytometry monocyte panel marker antibody panel. This antibody panel was used for both unfixed and fixed cell panels to identify cell-surface markers of monocyte subsets.

| **Table 3** |  |  |  |  |
| --- | --- | --- | --- | --- |
| Flow-cytometry monocyte marker antibody panel |  |  |  |  |
| **Target** | **Conjugate** | **Catalog #** | **Dilution** | **Company/Manufacturer** |
| CD14 | BV605 | BDB564054 | 1:25 | BD Biosciences |
| CD16 | BUV395 | BDB563785 | 1:50 | BD Biosciences |
| HLA-DR | PE-vio770 | 130-113-403 | 1:100 | Miltenyi Biotec |
| FcR block | - | 422301 | 1:20 | BioLegend |
| Live/dead Violet | - | L34962 | 1:2000 | Invitrogen |

### Fixed Cell Flow Cytometry and Staining for Lysosomal Health and LRRK2 Activity

After the 72-hour stimulation, cells were harvested and centrifuged at 300 x *g* for 5 min at 4°C. Supernatant was collected to quantify cytokine secretion (described below). Cell pellets were gently resuspended in 200 μL of cold PBS and transferred to a v-bottom 96-well plate (Sigma, CLS3896-48EA). Samples were centrifuged at 300 x *g* for 5 min at 4°C. Cells were resuspended in 200 μL of complete growth media, and LysoTracker™ Red DND-99 (Invitrogen, L7528) was added to reach a final concentration of 500 nM for T cells or 200 nM for monocytes. Cells were incubated for 1 hour at 37°C in the dark. Samples were centrifuged at 300 x *g* for 5 min at 4°C. Cell pellets were resuspended in PBS and washed x 2 by centrifugation at 300 x *g* for 5 min at 4°C. Cells were resuspended in 50 μL of Live/Dead Fixable Violet stain (diluted 1:2000 in PBS, Invitrogen, L34962) and incubated in the dark at room temperature for 30 min. Cells were centrifuged at 300 x *g* for 5 min at 4°C washed in PBS x 2. Cells were resuspended in 50 μL of of PBS containing diluted antibodies (see Table 2 for T cell panel, see Table 3 for monocyte panel) and incubated in the dark at 4°C for 20 min. Cells were centrifuged at 300 x *g* for 5 min at 4°C and washed x 2 in PBS. Cells were re-suspended and fixed in 100 μL of 1% paraformaldehyde (PFA) at 4°C in the dark for 30 min. Cells were washed 2 x with PBS, then resuspended in 100 μL of permeabilization buffer (eBiosciences, 00-8333-56) and incubated on ice for 15 min. Anti-pT73 Rab10 antibody (Abcam, ab241060) was added to each well at 0.55 μg per well and incubated at room temperature and protected from light for 30 min. Cells were centrifuged at 300 x *g* for 5 min at 4°C washed in PBS x 2. Cells were resuspended in 100 μL of PBS containing 1% normal goat/donkey serum, 2% BSA and 1:1000 AF488 donkey anti-rabbit secondary (Thermo Fisher, A-21206) and incubated at room temperature and protected from light for 30 min. Cells were centrifuged at 300 x *g* for 5 min at 4°C washed in PBS x 2. Cells were resuspended in 100 μL of PBS containing 1% normal goat/donkey serum, 2% BSA 1:100 anti-LRRK2 AF700 antibody and incubated at 4°C covered for 20 min. Cells were centrifuged at 300 x *g* for 5 min at 4°C, and then washed in FACS buffer × 3. Cells were analyzed via flow cytometry on a FACSymphony™ A3 cytometer (BD Biosciences). Data were analyzed using FlowJo version 10.10.0 software.

### Cytokine Quantification

V-PLEX custom Human Biomarkers kit (Meso Scale Discovery (MSD), K151ARH-2) was used to quantify cytokines in conditioned media from cultured T cells and monocytes. Media was diluted 1:4 with MSD kit diluent and incubated in duplicate at room temperature in the provided MSD plate with capture antibodies for 2 hours as per manufacturer’s instructions. Plates were then washed x 3 with PBS with 0.05% Tween-20 and detection antibodies conjugated with electrochemiluminescent labels were added and incubated at room temperature for another 2 hours. After 3 x washes with PBS containing 0.05% Tween-20, MSD buffer was added and the plates were loaded into the QuickPlex MSD instrument for quantification.

### Statistics

Data and statistical analyses were performed using GraphPad Prism 10. For assessing differences between groups, data were analyzed by either one-way or two-way analysis of variance (ANOVA), or by *t*-test. In instances when data did not fit parametric assumptions, Kruskal–Wallis non-parametric ANOVA was used. *Post-hoc* tests following ANOVAs were conducted using Tukey’s method for correcting for multiple comparisons. For assessing relationships between read-outs, data were analyzed by Pearson’s *r*. In instances when data did not fit parametric assumptions, Spearman’s rank was used to assess relationships between variables. Two-tailed levels of significance were used and *p* < 0.05 was considered statistically significant. Graphs are depicted by means ± standard error of the mean (SEM).

## Acknowledgements

We thank the participants in this study for their support of our research, as well as the movement disorder specialists and clinical research coordinators for their tireless work recruiting subjects and collecting the samples needed for this study, in particular Julie Segura. We thank the members of the Tansey lab for useful discussions. This work was primarily supported by funds from the Michael J. Fox Foundation under the MJFF-019562 award to M.G.T and R.L.W. This work was partially supported by the National Institutes of Health (R25AG076396). We also thank the McKnight Brain Institute, the Normal Fixel Institute for Neurological Diseases, and the Center for Translational Research in Neurodegenerative Disease at the University of Florida. We thank all the members of the Tansey research team, and the University of Florida ICBR Cytometry Core Facility (RRID:SCR_019119) for their assistance. Some schematics (Fig. 1) were created with BioRender.com.

## Author Contributions

J.R.M. was responsible for experimental design, plotting and analyzing data, data interpretation, and drafting and editing the manuscript. J.R.M. and A.M.T. were responsible for performing experiments, optimizing experimental workflow, and optimizing experimental design. H.A.S. was responsible for running all MSD-experiments. S.M. and S.A. were responsible for sample collection. S.A. and N.R.M. were responsible for patient recruitment. M.G.T. and R.L.W. were responsible for conceiving the project, funding acquisition, supervision, and project administration. All authors participated in writing and editing of the manuscript.

