## Supplemental Data for "Peripheral immune cell response to stimulation stratifies Parkinson’s disease progression from prodromal to clinical stages"

### Supplementary Fig. 1: Monocyte absolute cytokine secretion

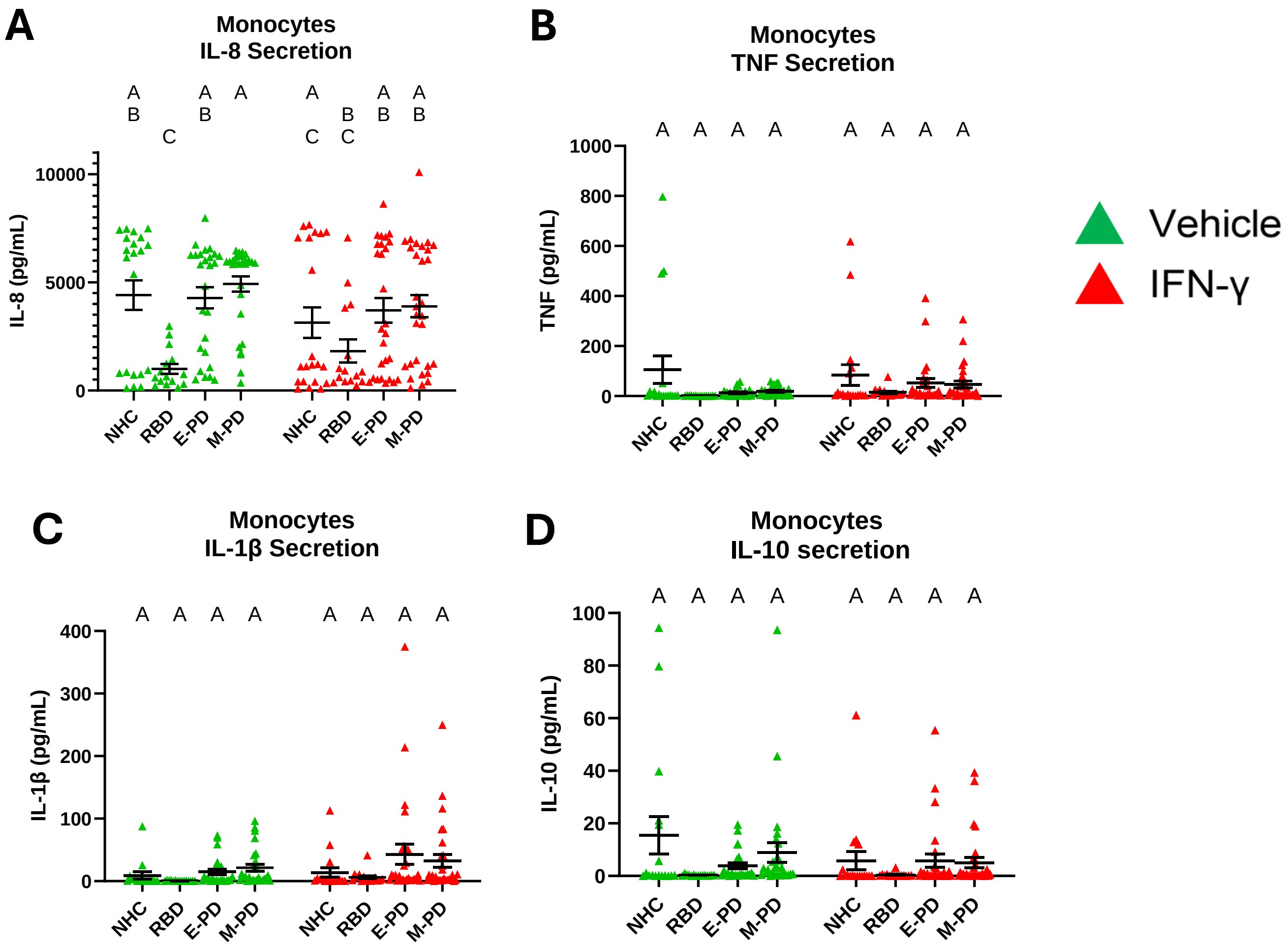


**Supplementary Fig. 1: Monocyte absolute cytokine secretion.** Bar graphs overlaid with scatter plots showing absolute concentrations of secreted inflammatory cytokines from isolated monocytes from NHCs, RBD patients, EPD patients, and MPD patients. Absolute concentrations of secreted TNF (**A**), IL-1β (**B**), IL-8 (**C**), and IL-10 (**D**). Bars represent mean +/- SEM. NHC neurologically healthy controls, $n$ = 21; RBD patients with REM sleep behavior disorder, $n$ = 15; EPD patients with early-stage PD, $n$ = 27; MPD patients with moderate-stage PD, $n$ = 30. Each symbol represents the measurement from a single individual. The results in **A-D** were analyzed using two-way ANOVA with Tukey's corrections for multiple comparisons. Groups sharing the same letters are not significantly different ($p>0.05$) whilst groups displaying different letters are significantly different ($p<0.05$).

### Supplementary Fig. 2: T cell absolute cytokine secretion

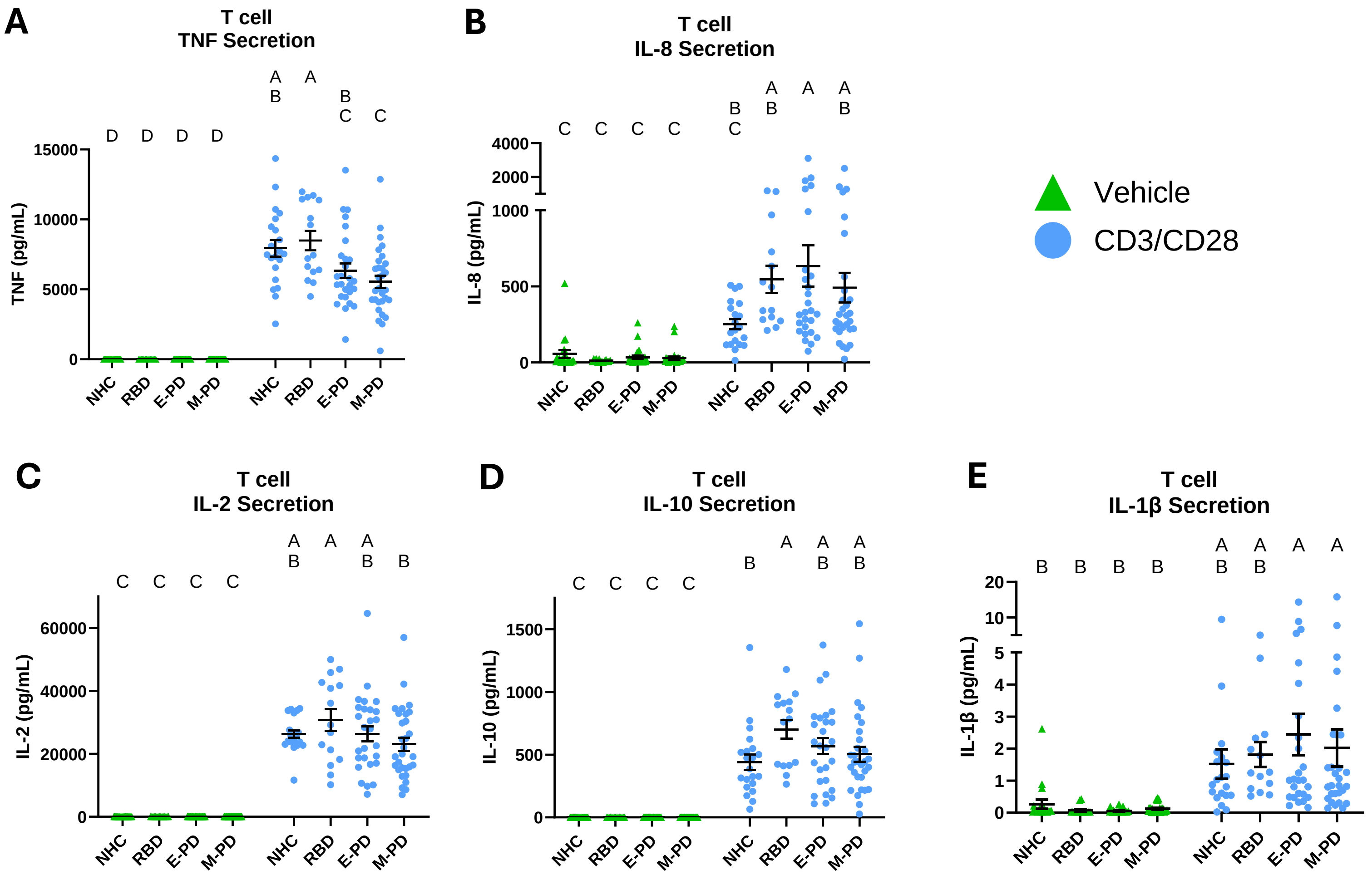


**Supplementary Fig. 2: T cell absolute cytokine secretion.** Bar graphs overlaid with scatter plots showing absolute concentrations of secreted inflammatory cytokines from isolated CD3+ T cells from NHCs, RBD patients, EPD patients, and MPD patients. Absolute concentrations of secreted TNF (**A**), IL-1β (**B**), IL-8 (**C**), and IL-10 (**D**). Bars represent mean +/- SEM. NHC neurologically healthy controls, *n* = 21; RBD patients with REM sleep behavior disorder, *n* = 15; EPD patients with early-stage PD, *n* = 27; MPD patients with moderate-stage PD, *n* = 30. Each symbol represents the measurement from a single individual. The results in **A-D** were analyzed using two-way ANOVA with Tukey's corrections for multiple comparisons. Groups sharing the same letters are not significantly different (p>0.05) whilst groups displaying different letters are significantly different (p<0.05).

Supplementary Fig. 3: Flow cytometry gating strategy for T cells using fluorescence-minus-one controls

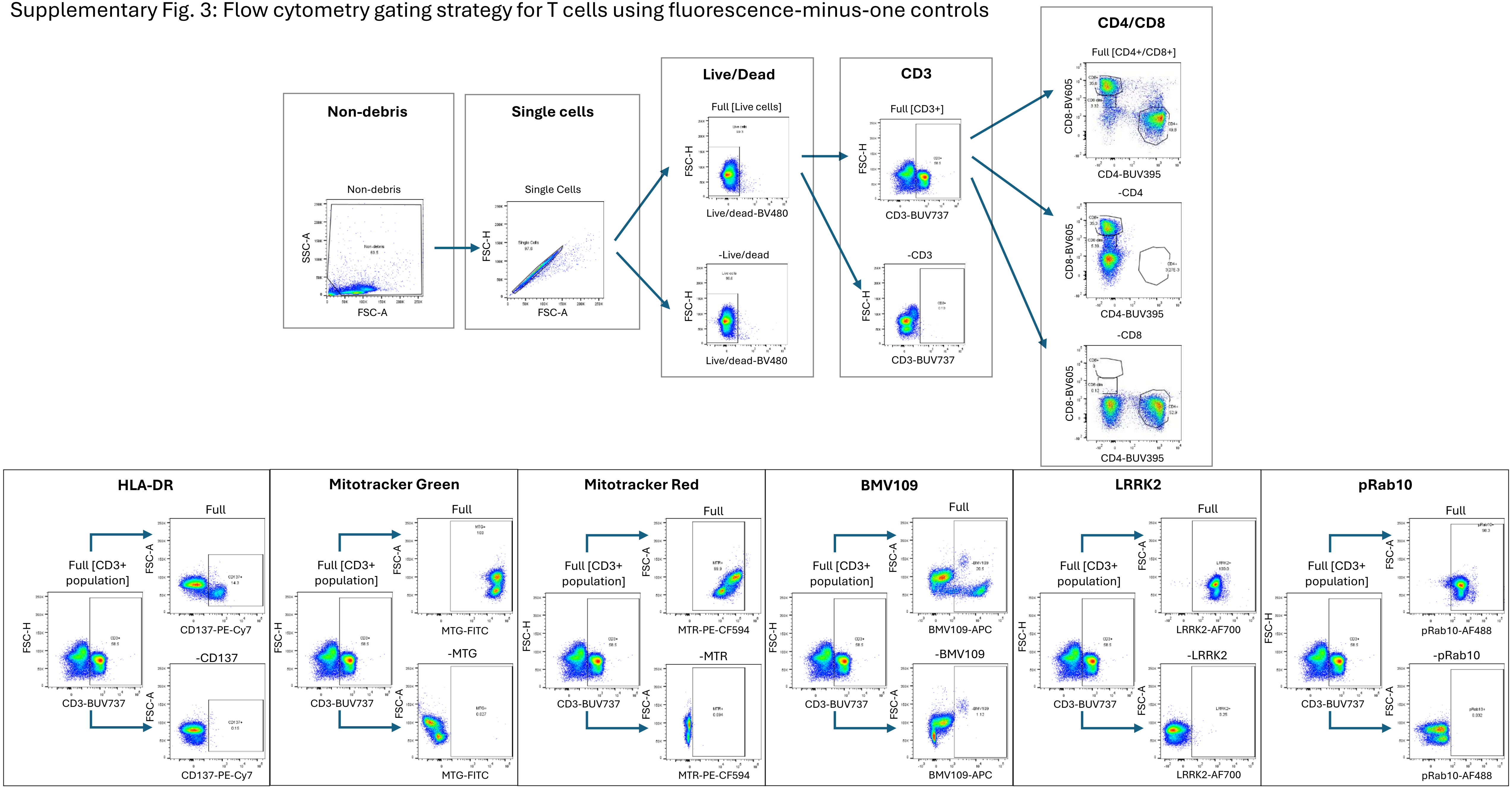


**Supplementary Fig. 3: Flow cytometry gating strategy for T cells using fluorescence-minus-one controls**. Total PBMCs were stained with antibody-fluorophore conjugates and gates were defined for positive and negative populations based on <1% negatively stained cells in FMOC.

Supplementary Fig. 4: Flow cytometry gating strategy for monocytes using fluorescence-minus-one controls

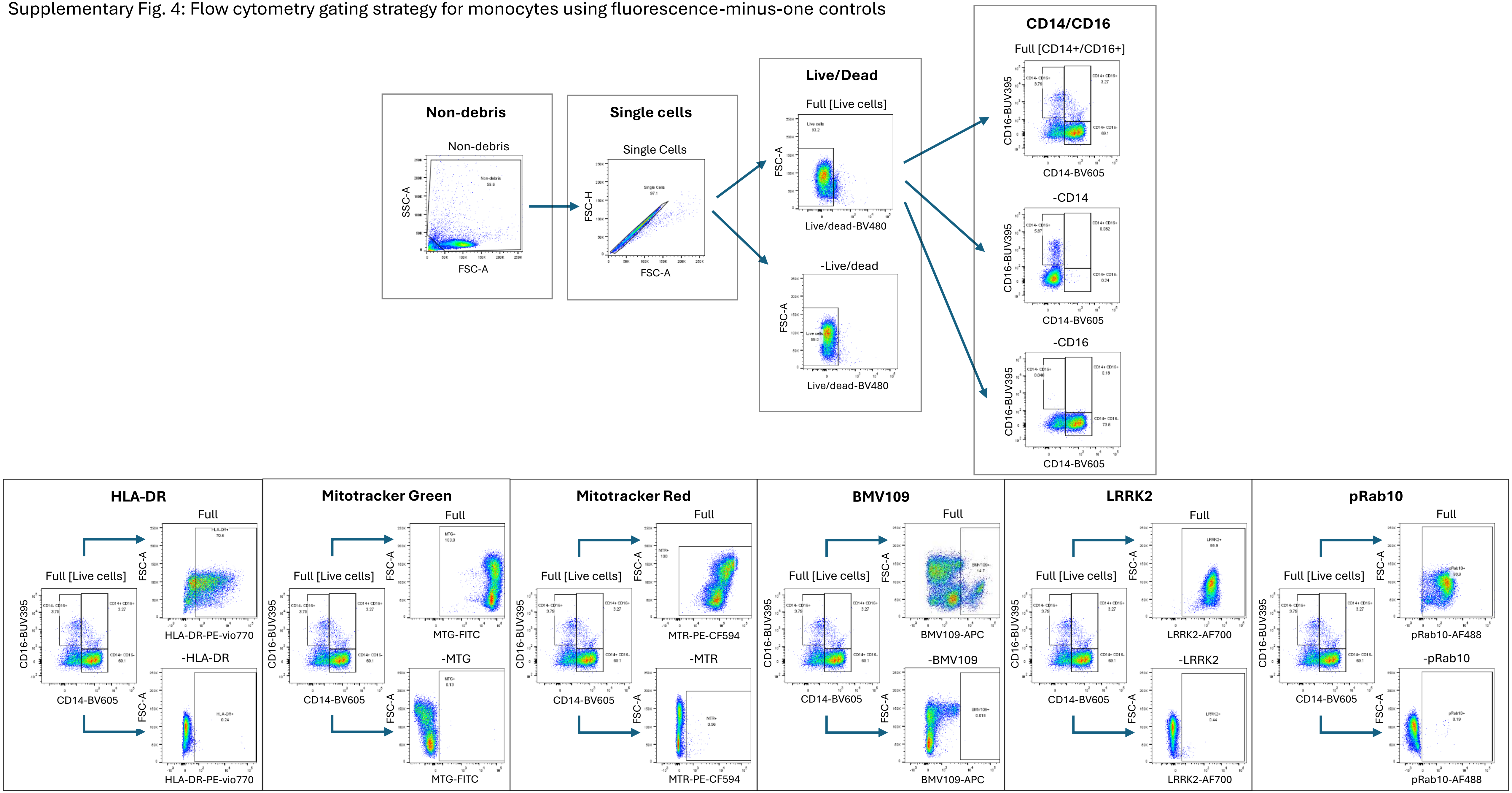


**Supplementary Fig. 4: Flow cytometry gating strategy for monocytes using fluorescence-minus-one controls**. Isolated monocytes were stained with antibody-fluorophore conjugates and gates were defined for positive and negative populations based on <1% negatively stained cells in FMOC.

### Supplementary Fig. 5: PBMC subtype counts

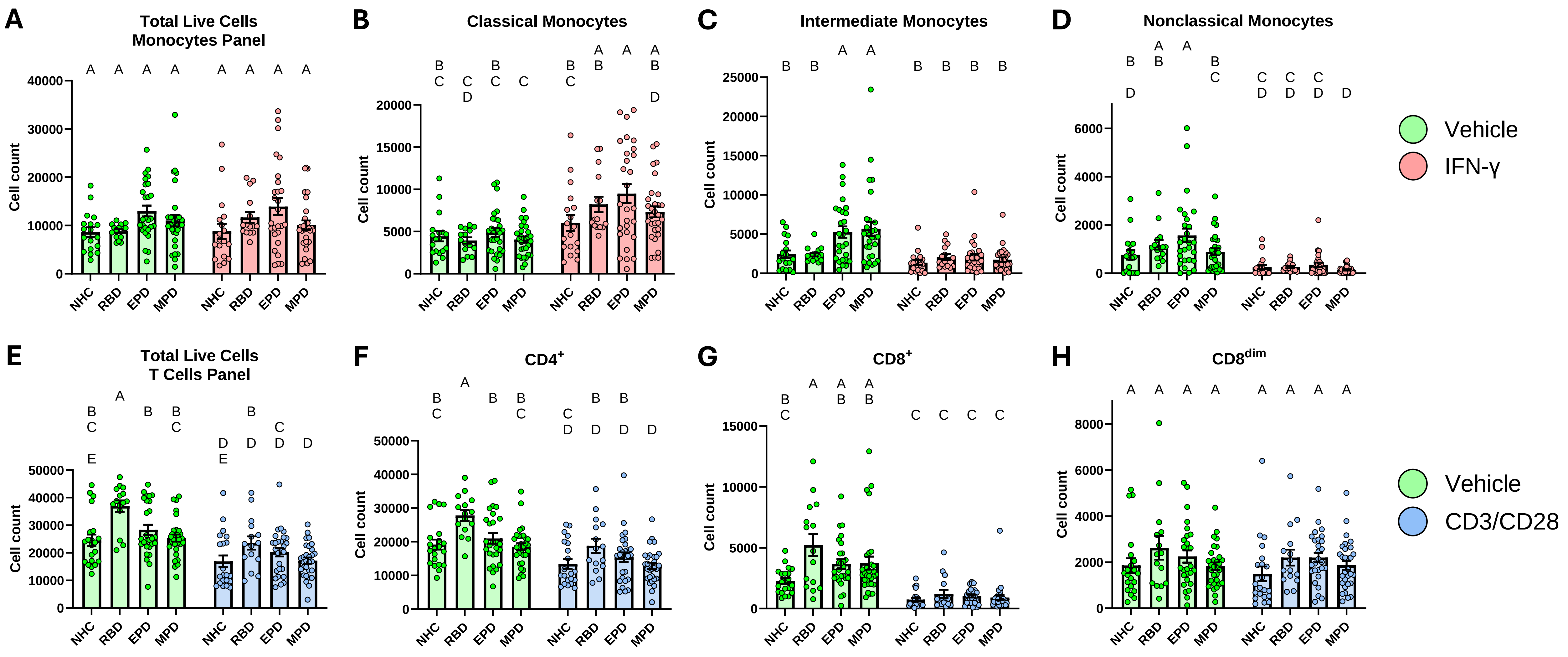


**Supplementary Fig. 5: PBMC subtype counts**. Bar graphs overlaid with scatter plots showing the raw counts of monocytes and T cells in PBMCs from NHCs, RBD patients, EPD patients, and MPD patients. **A** Raw counts of total monocytes ($CD14^{+}$ and/or $CD16^{+}$) among live cells. **B** Raw counts of classical monocytes ($CD14^{+}CD16^{-}$) among total monocytes. **C** Raw counts of intermediate monocytes ($CD14^{+}CD16^{+}$) among total monocytes. **D** Raw counts of nonclassical monocytes ($CD14^{dim}CD16^{+}$) among total monocytes. **E** Raw counts of total $CD3^{+}$ T cells among total live cells. **F** Raw counts of $CD4^{+}CD8^{-}$ among total $CD3^{+}$ T lymphocytes. **G** Raw counts of $CD4^{-}CD8^{+}$ among total $CD3^{+}$ T lymphocytes. **H** Raw counts of $CD4^{-}CD8^{dim}$ among total $CD3^{+}$ T lymphocytes. Bars represent mean +/- SEM. NHC neurologically healthy controls, *n* = 21; RBD patients with REM sleep behavior disorder, *n* = 15; EPD patients with early-stage PD, *n* = 27; MPD patients with moderate-stage PD, *n* = 30. Each symbol represents the measurement from a single individual. The results in **A-H** were analyzed using two-way ANOVA with Tukey's corrections for multiple comparisons. Groups sharing the same letters are not significantly different (p>0.05) whilst groups displaying different letters are significantly different (p<0.05). Samples were not run to completion through flow cytometer, therefore these counts do not represent the total number of cells collected.

### Supplementary Fig. 6: Monocyte mitochondrial health across multiple stages of PD

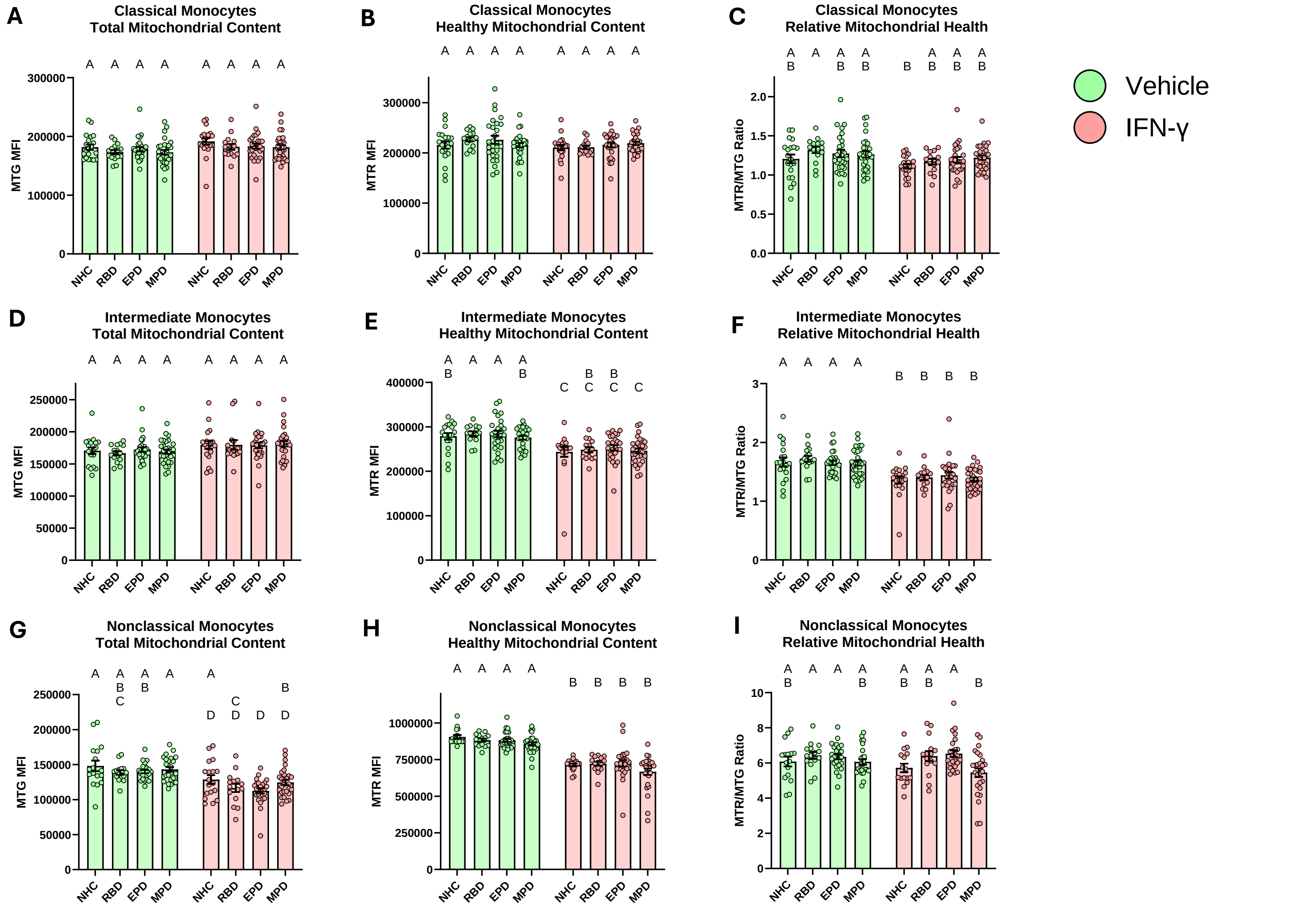


**Supplementary Fig. 6: Monocyte mitochondrial health across multiple stages of PD**. Bar graphs overlaid with scatter plots showing the mitochondrial content and mitochondrial health after immune stimulation of monocyte subsets from NHCs, RBD patients, EPD patients, and MPD patients. **A** Total mitochondrial content of classical monocytes (CD14$^+$CD16$^-$). **B** Healthy mitochondrial content with negative membrane potential in classical monocytes (CD14$^+$CD16$^-$). **C** Ratio of healthy mitochondrial content divided by the total in classical monocytes (CD14$^+$CD16$^-$). **D** Total mitochondrial content of intermediate monocytes (CD14$^+$CD16$^+$). **E** Healthy mitochondrial content with negative membrane potential in intermediate monocytes (CD14$^+$CD16$^+$). **F** Ratio of healthy mitochondrial content divided by the total in intermediate monocytes (CD14$^+$CD16$^+$). **G** Total mitochondrial content of nonclassical monocytes (CD14$^{dim}$CD16$^+$). **H** Healthy mitochondrial content with negative membrane potential in nonclassical monocytes (CD14$^{dim}$CD16$^+$). **I** Ratio of healthy mitochondrial content divided by the total in nonclassical monocytes (CD14$^{dim}$CD16$^+$). Bars represent mean +/- SEM. NHC neurologically healthy controls, *n* = 21; RBD patients with REM sleep behavior disorder, *n* = 15; EPD patients with early-stage PD, *n* = 27; MPD patients with moderate-stage PD, *n* = 30. Each symbol represents the measurement from a single individual. The results in **A-I** were analyzed using two-way ANOVA with Tukey's corrections for multiple comparisons. Groups sharing the same letters are not significantly different (p>0.05) whilst groups displaying different letters are significantly different (p<0.05). MTG MitoTracker Green FM, MTR MitoTracker Red CMXRos.

### Supplementary Fig. 7: Motor severity compared to stimulation-dependent cytokine secretion

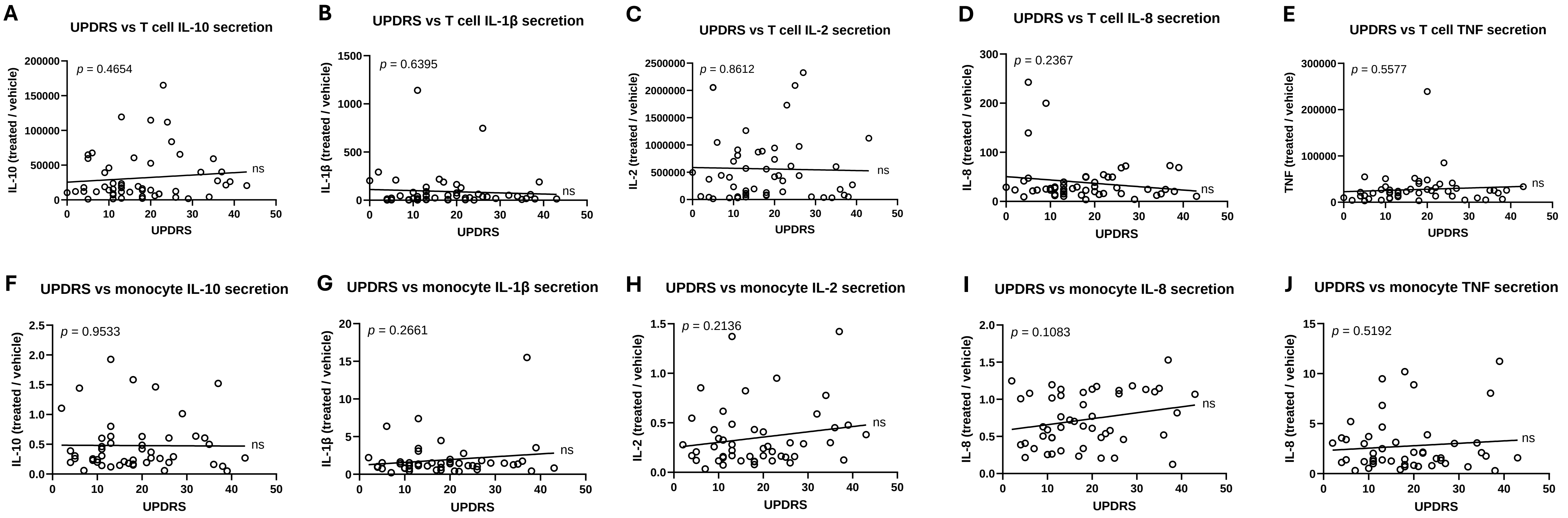


**Supplementary Fig. 7: Motor severity compared to stimulation-dependent cytokine secretion**. Correlational analysis showing the relationship between UPDRS scores from Parkinson's disease (PD) patients and stimulation-dependent cytokine secretion from peripheral immune cell subsets. Correlation between UPDRS scores from early and moderate-stage Parkinson's disease patients and stimulation-dependent secretion of IL-10 (**A**), IL-1β (**B**), IL-2 (**C**), IL-8 (**D**), and TNF (**E**) from isolated T cells. Correlation between UPDRS scores from early and moderate-stage Parkinson's disease patients and stimulation-dependent secretion of IL-10 (**F**), IL-1β (**G**), IL-2 (**H**), IL-8 (**I**), and TNF (**J**) from isolated monocytes. *p*-value based on Pearson correlation. early-stage PD, $n = 23$; moderate-stage PD, $n = 27$. Each symbol represents the measurement from a single individual. * signifies that the slope of the line is significantly different from zero ($p < 0.05$). *ns* signifies that the slope of the line is not significantly different from zero ($p > 0.05$).

### Supplementary Fig. 8: Lysosomal content of monocyte and T cell subsets across multiple stages of PD

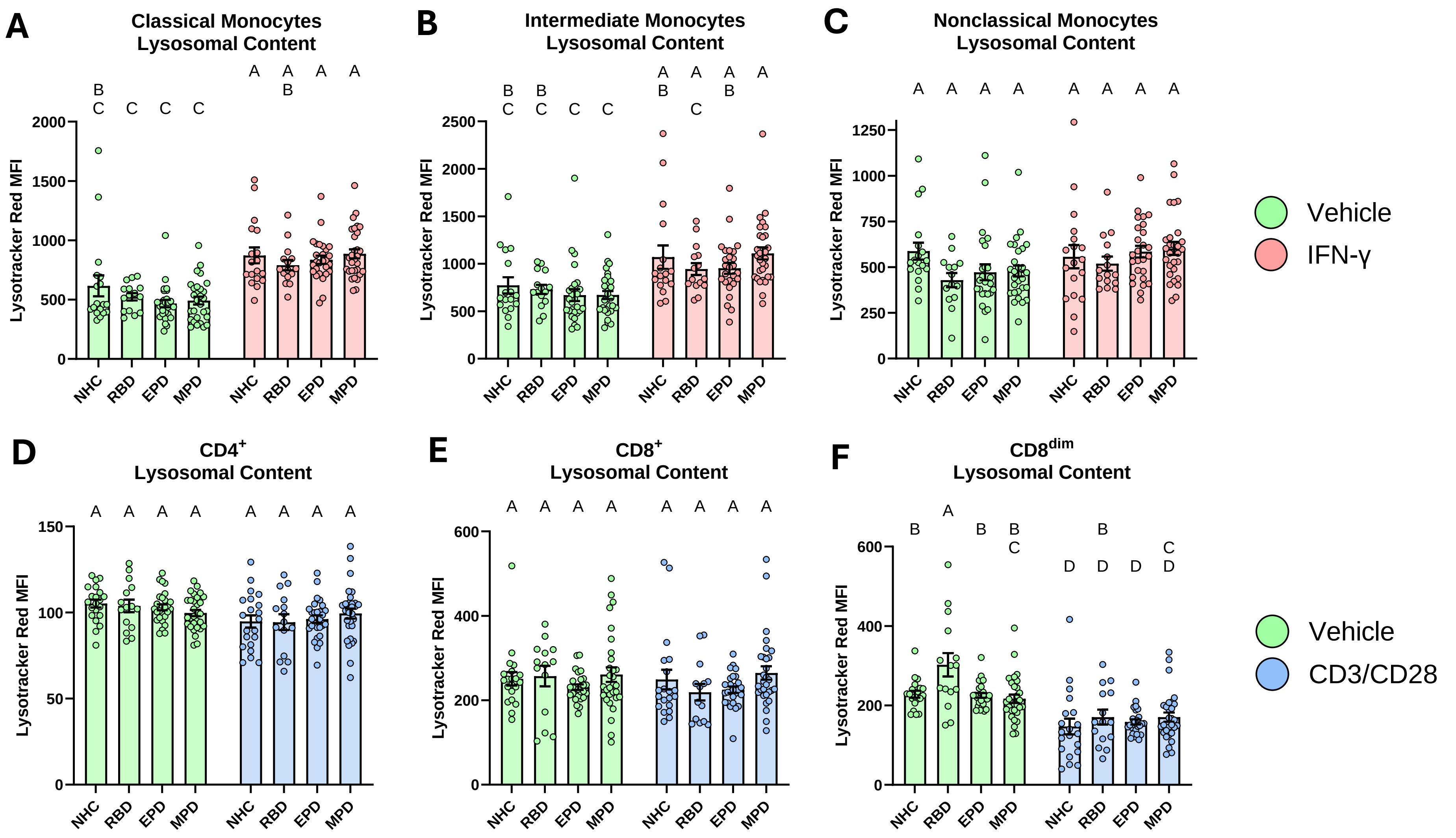


**Supplementary Fig. 8: Lysosomal content of monocyte and T cell subsets across multiple stages of PD**. Bar graphs overlaid with scatter plots showing the lysosomal content after immune stimulation of monocyte and T cell subsets from NHCs, RBD patients, EPD patients, and MPD patients. Lysosomal content was quantified using MFI of Lysotracker Red DND. **A** Total lysosomal content of classical monocytes ($CD14^{+}CD16^{-}$). **B** Total lysosomal content of intermediate monocytes ($CD14^{+}CD16^{+}$). **C** Total lysosomal content of nonclassical monocytes ($CD14^{dim}CD16^{+}$). **D** Total lysosomal content of $CD4^{+}/CD8^{-}$ T cells. **E** Total lysosomal content of $CD4^{-}/CD8^{+}$ T cells. **H** Total lysosomal content of $CD4^{-}/CD8^{dim}$ T cells. Bars represent mean +/- SEM. NHC neurologically healthy controls, *n* = 21; RBD patients with REM sleep behavior disorder, *n* = 15; EPD patients with early-stage PD, *n* = 27; MPD patients with moderate-stage PD, *n* = 30. Each symbol represents the measurement from a single individual. The results in **A-F** were analyzed using two-way ANOVA with Tukey's corrections for multiple comparisons. Groups sharing the same letters are not significantly different (p>0.05) whilst groups displaying different letters are significantly different (p<0.05). MFI median fluorescence intensity.

### Supplementary Fig. 9: Pan-cathepsin activity of T cell subsets

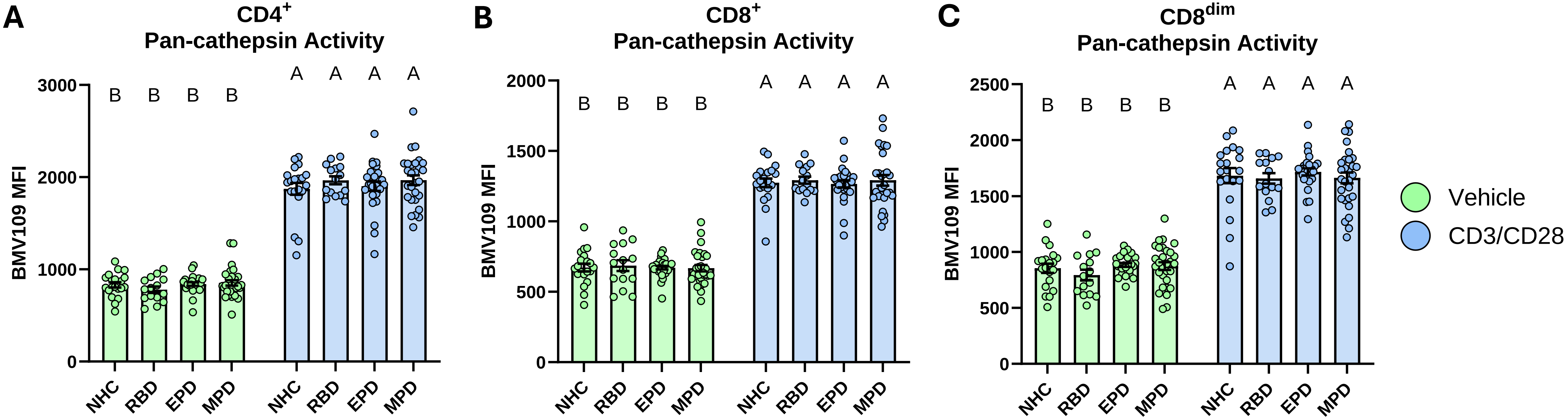


**Supplementary Fig. 9: Pan-cathepsin activity in T cell subsets**. Bar graphs overlaid with scatter plots showing the pan-cathepsin activity after CD3/CD28 Dynabead stimulation of T cell subsets from NHCs, RBD patients, EPD patients, and MPD patients. Pan-cathepsin activity was quantified using MFI of BMV109. **A** Pan-cathepsin activity of $CD4^+/CD8^-$ T cells. **B** Pan-cathepsin activity of $CD4^-/CD8^+$ T cells. **C** Pan-cathepsin activity of $CD4^-/CD8^{dim}$ T cells. Bars represent mean +/- SEM. NHC neurologically healthy controls, *n* = 21; RBD patients with REM sleep behavior disorder, *n* = 15; EPD patients with early-stage PD, *n* = 27; MPD patients with moderate-stage PD, *n* = 30. Each symbol represents the measurement from a single individual. The results in **A-C** were analyzed using two-way ANOVA with Tukey's corrections for multiple comparisons. Groups sharing the same letters are not significantly different ($p>0.05$) whilst groups displaying different letters are significantly different ($p<0.05$). MFI median fluorescence intensity.

### Supplementary Fig. 10: LRRK2 expression and pRab10 expression in monocyte subsets

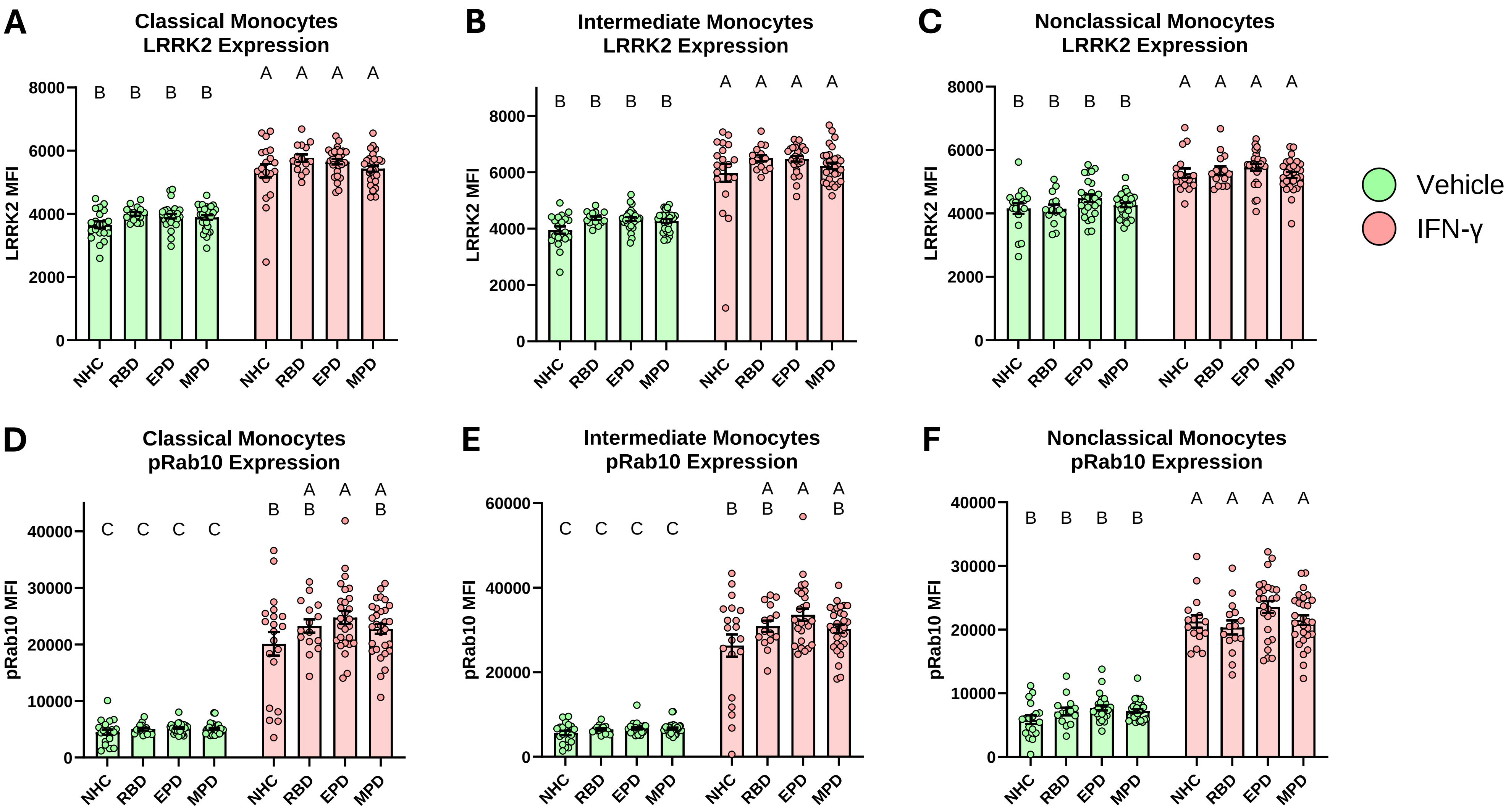


**Supplementary Fig. 10: LRRK2 expression and pRab10 expression in monocyte subsets**. Bar graphs overlaid with scatter plots showing the expression of LRRK2 and pRab10 after IFNγ stimulation of monocyte subsets from NHCs, RBD patients, EPD patients, and MPD patients. **A** LRRK2 expression in classical monocytes ($CD14^{+}CD16^{-}$). **B** LRRK2 expression in intermediate monocytes ($CD14^{+}CD16^{+}$). **C** LRRK2 expression in nonclassical monocytes ($CD14^{dim}CD16^{+}$). **D** pRab10 expression in classical monocytes ($CD14^{+}CD16^{-}$). **E** pRab10 expression in intermediate monocytes ($CD14^{+}CD16^{+}$). **F** pRab10 expression in nonclassical monocytes ($CD14^{dim}CD16^{+}$). Bars represent mean +/- SEM. NHC neurologically healthy controls, $n$ = 21; RBD patients with REM sleep behavior disorder, $n$ = 15; EPD patients with early-stage PD, $n$ = 27; MPD patients with moderate-stage PD, $n$ = 30. Each symbol represents the measurement from a single individual. The results in **A-F** were analyzed using two-way ANOVA with Tukey's corrections for multiple comparisons. Groups sharing the same letters are not significantly different ($p>0.05$) whilst groups displaying different letters are significantly different ($p<0.05$). LRRK2 leucine-rich repeat kinase 2, pRab10 phosphorylated Rab10.

### Supplementary Fig. 11: LRRK2 expression and pRab10 expression in T cell subsets

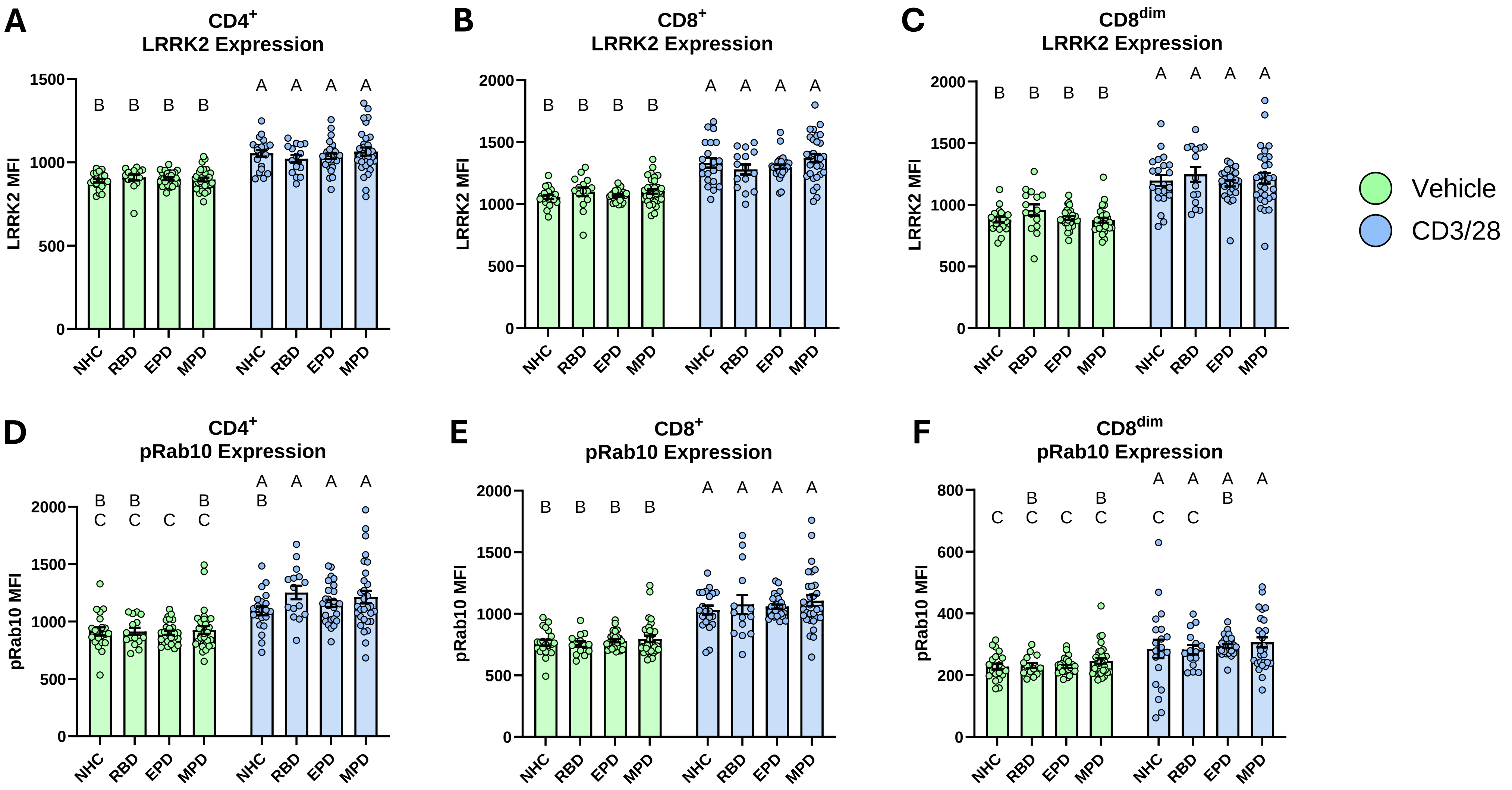


**Supplementary Fig. 11: LRRK2 expression and pRab10 expression in T cell subsets**. Bar graphs overlaid with scatter plots showing the expression of LRRK2 and pRab10 after CD3/CD28 stimulation of T cell subsets from NHCs, RBD patients, EPD patients, and MPD patients. **A** LRRK2 expression in $CD4^+/CD8^-$ T cells. **B** LRRK2 expression in $CD4^-/CD8^+$ T cells. **C** LRRK2 expression in $CD4^-/CD8^{dim}$ T cells. **D** pRab10 expression in $CD4^+/CD8^-$ T cells. **E** pRab10 expression in $CD4^-/CD8^+$ T cells. **F** pRab10 expression in $CD4^-/CD8^{dim}$ T cells. Bars represent mean +/- SEM. NHC neurologically healthy controls, $n$ = 21; RBD patients with REM sleep behavior disorder, $n$ = 15; EPD patients with early-stage PD, $n$ = 27; MPD patients with moderate-stage PD, $n$ = 30. Each symbol represents the measurement from a single individual. The results in **A-F** were analyzed using two-way ANOVA with Tukey's corrections for multiple comparisons. Groups sharing the same letters are not significantly different ($p>0.05$) whilst groups displaying different letters are significantly different ($p<0.05$). LRRK2 leucine-rich repeat kinase 2, pRab10 phosphorylated Rab10.
